# The EXO70B2 exocyst subunit contributes to papillae and encasement formation in anti-fungal defence in Arabidopsis

**DOI:** 10.1101/2021.06.20.449050

**Authors:** Jitka Ortmannová, Juraj Sekereš, Ivan Kulich, Jiří Šantrůček, Petre Dobrev, Viktor Žárský, Tamara Pečenková

**Author notes:** Correspondence: Tamara Pečenková. authors email addresses.

## Abstract

In the reaction to non-adapted *Blumeria graminis* f. sp. *hordei* (Bg), *Arabidopsis thaliana* leaf epidermal cells deposit cell wall reinforcements called papillae or seal fungal haustoria in encasements, both of which involve intensive exocytosis. A plant syntaxin SYP121/PEN1 has been found to be of key importance for the timely formation of papillae, and the vesicle tethering complex exocyst subunit EXO70B2 has been found to contribute to their morphology. Here, we identify a specific role for the EXO70B2-containing exocyst complex in the papillae membrane domains important for the callose deposition and GFP-SYP121 delivery to the focal attack sites, as well as its contribution to encasement formation. The mRuby2-EXO70B2 co-localises with the exocyst core subunit SEC6 and GFP-SYP121 in the membrane domain of papillae, and both proteins have the capacity to directly interact. The *exo70B2/syp121* double mutant has a reduced number of papillae and haustorial encasements in response to Bg, indicating an additive role of the exocyst in SYP121 coordinated non-host resistance. In summary, we report cooperation between the plant exocyst and a SNARE protein in penetration resistance against non-adapted fungal pathogens.

**Highlight:** The exocyst complex containing EXO70B2 subunit has specific role in papilla and encasement formation, implemented in coordination with the pathway regulated by the SYP121 SNARE complex in defence.

## Introduction

One of the main tiers of plant non-host resistance is a secretory pathway. At the moment of detection of a pathogen at the plant surface, pathogen-associated molecular pattern (PAMP)-triggered immunity is activated and plant cells start to react and re-polarize their secretory pathway to the attack sites (Schmelzer, 2002; Jones and Dangl, 2006; Lee *et al.*, 2017). The newly established secretory domain allows the plant cells to deposit focal cell wall reinforcements called papillae (Bestwick *et al.*, 1995). Their composition is a mixture of cell wall and antimicrobial components, such as callose (beta-1,3-glucan), pectins, lignins, reactive oxygen species (ROS), phytoalexins, and thionins (Schmelzer, 2002; Chowdhury *et al.*, 2014). Despite the surface barriers, some fungal hyphae may invade plant cells with feeding structures called haustoria. Haustoria are surrounded by a specialized extrahaustorial membrane, the origin of which remains elusive (EHM) (Micali *et al.*, 2011). To stop fungal growth, plant cells seal the haustoria by encasements, defensive structures made of similar components to the papillae (Heath and Heath, 1971; Zeyen *et al.*, 2002).

In Arabidopsis thaliana, non-host resistance remains sufficient against evolutionarily non-adapted fungi, such as *Erysiphe pisi* (Ep) or Bg (Vogel and Somerville, 2000; Lipka *et al.*, 2005; Consonni *et al.*, 2006). A forward genetic screen revealed that the PENETRATION1 (PEN1/SYP121) locus plays a crucial role in this non-host resistance (Collins *et al.*, 2003; Stein *et al.*, 2006; Lipka *et al.*, 2008). SYP121 belongs to the family of plasma membrane (PM) Qa-Soluble N-ethylmaleimide-sensitive factor Attachment protein REceptors (SNAREs). SYP121 forms a ternary complex with the cytoplasmic Qbc-SNARE SNAP33 (soluble N-ethylmaleimide-sensitive factor adaptor protein 33) and the R-SNAREs VAMP721 or VAMP722 (vesicle-associated membrane protein 721/722) (Kwon *et al.*, 2008). This complex promotes the exocytosis of secretory vesicles carrying defence-related cargoes to plant cell/fungus contact sites and mediates papillae formation (Assaad *et al.*, 2004; Kwon *et al.*, 2008; Meyer *et al.*, 2009). GFP-SYP121 also associates in a GNOM-dependent manner with exosomes in the extracellular matrix of both papillae and encasements (Nielsen *et al.*, 2012; Rutter and Innes, 2017). However, the canonical role of SYP121 probably relates to intracellular endomembranes fusions, possibly at the Golgi (Nielsen and Thordal-Christensen, 2012, 2013; Nielsen *et al.*, 2017). The SYP121-interacting R-SNARE VAMP727 has been found to operate on Rab-GTPase ARA7-positive multivesicular bodies (MVB), indicative of the involvement of these MVB in secretion into the papillary matrix (An *et al.*, 2007; Ebine *et al.*, 2011; Nielsen *et al.*, 2012; Heard *et al.*, 2015).

In yeasts and mammals, SNARE proteins execute their function in cooperation with other proteins and protein complexes, one of which is the exocyst complex (Hsu *et al.*, 1996; Sivaram *et al.*, 2005; Dubuke *et al.*, 2015; Yue *et al.*, 2017). The exocyst vesicle tethering complex assembles from the core subunits SEC3 (membrane lipid-interacting subunits), SEC5, SEC6, SEC8, SEC10, SEC15 and EXO84, and the plasma membrane lipid-interacting and localizing subunit EXO70 (TerBush *et al.*, 1996; Guo *et al.*, 1997; Liu *et al.*, 2018; Zhu *et al.*, 2019). While being encoded by a single gene in the yeast and mammalian genome, EXO70 is encoded by many paralogs in angiosperm genomes (Elias *et al.*, 2003; Cvrcková *et al.*, 2012). In accordance, the plant exocyst complex has multiple functions, e. g. seed coat deposition, tip growth, cytokinesis, recycling and autophagy (Cole *et al.*, 2005; Synek *et al.*, 2006; Hála *et al.*, 2008; Fendrych *et al.*, 2010, 2013; Kulich *et al.*, 2010; Drdova *et al.*, 2013). It has been hypothesized that the different isoforms may target the exocyst and exocytosis to different plant cell PM domains even within a single cell type such as a growing pollen tube or trichomes (Žárskỳ *et al.*, 2009, 2013; Sekereš *et al.*, 2017; Synek *et al.*, 2017; Kubátová *et al.*, 2019).

In Arabidopsis, the specific role of the EXO70 isoform EXO70B2 has been documented in PAMP-triggered immunity regulation and papillae biogenesis (Pečenková *et al.*, 2011; Stegmann *et al.*, 2012; Wang *et al.*, 2020). Here, we show that in addition to EXO70B2-GFP, the core exocyst subunit SEC6 tagged with GFP is also localised to papillae and encasement structures indicating that whole exocyst complex is there. Further on, we observed an increase in the penetration rate of the non-adapted powdery mildew fungi Bg (and also Ep) in two exocyst core subunit mutants as a probable consequence of slower callose deposition. The EXO70B2 and SYP121 have a capacity to directly interact and to mutually affect papilla-related localisations. The encasements formation and callose accumulation are reduced in the double mutant *exo70B2*/*syp121*. These results indicate that the fully functional exocyst complex involving the isoform EXO70B2 is important for penetration resistance, likely as a component of the previously described SYP121-dependent secretory pathway.

## Materials and methods

### Plant material

The seeds of Arabidopsis thaliana were sterilised and plated on 1/2 MS with vitamins, 1% sucrose medium. The plants were grown in vitro for 10 days and used for qRT-PCR, confocal imaging, or transferred to Jiffy tablets and grown in a growth chamber under short day conditions (21°C, 10/14 h light/dark, 80% humidity and light intensity of 125 μmol m−2 s−1 in a range from 400–700 nm).

The mutant lines of Arabidopsis used in this study were as follows: knockout line (KO) *sec5a-1* (GABI_731C01, Fig. S6), KO line *syp121-1* (Collins *et al.*, 2003), knock-down line *sec8-m4* (Cole *et al.*, 2005), *exo70B2-2* KO line (Pečenková *et al.*, 2011). The transgenic lines of Arabidopsis that were used in this study were: EXO70B2: EXO70B2-GFP; EXO70B2:GFP-EXO70B2 (this work); SYP121:GFP-SYP121 (Kato *et al.*, 2010); SEC6:SEC6-GFP and 35S:GFP-EXO70A1 (Fendrych *et al.*, 2010); EXO70B1:GFP-EXO70B1 (Kulich *et al.*, 2013).

### Plasmid construction and generation of transgenic lines

All constructs (GFP-EXO70B2, EXO70B2-GFP and mRuby2-EXO70B2) were prepared with the Phusion PC (NEB, USA) reaction using as a template either genomic DNA for intron-less genes and promoters or cDNA. A list of the primers used is presented in Supplementary Table 1, as well as the lengths of the amplified fragments, restriction sites used for the cloning procedure and destination vectors used in the cloning procedure. The sequenced vectors were used to transform the *Agrobacterium tumefaciens* GV3101 strain. Arabidopsis WT or respective mutants were transformed by the Agrobacterium-mediated floral dip method (Clough and Bent, 1998).

### Pathogen inoculation and cytology

For the pathogen inoculation experiments, plants were cultivated under short day conditions (21°C, 10/14 h, 75% humidity with a light intensity of 125 μmol m−2 s−1). Bg was cultivated continuously on fresh barley (Golden promise) grown under short day conditions (19°C, 10/14 h, 50% humidity, and a light intensity of 70 μmol m−2 s−1). Ep was cultivated continuously on fresh pea (*Pisum sativum* variety Petit provencal) under short day conditions (19°C, 10/14 h, 60% humidity and light intensity of 70 μmol m−2 s−1). The *Blumeria graminis f. sp. hordei* isolate A6 on barley genotype P01 was kindly provided by Lenka Burketová, Laboratory of Pathological Plant Physiology in Prague, IEB AS, Czech Republic. Paul Schulze-Lefert from the Max Planck Institute for Plant Breeding Research, Cologne, kindly provided the *Erysiphe pisi* isolate. The plants, approximately 4 weeks old, were inoculated by spreading spores from infected barley or pea onto the adaxial site of their leaves (from leaf to leaf). The 5th and 6th leaves were cut off at selected hpi and cleared with 96% ethanol or chloral hydrate. For callose visualisation, cleared leaves were stained with an alkali solution of aniline blue (Eschrich and Currier, 1964). For penetration rate visualisation, fungal structures were stained with 250 mg/ml trypan blue in lactophenol/ethanol solution (Vogel and Somerville, 2000). For quantification purposes, we considered only epidermal cells that were attacked by a single germinated spore. Stained leaves were observed with classical epifluorescence microscopy or optical microscopy using a Nikon Eclipse TE 2000-E inverted microscope. For the bacterial sensitivity assay, the Arabidopsis seedling flood-inoculation assay with *Pseudomonas syringae pv. tomato* DC3000 was utilized according to (Ishiga *et al.*, 2011) with minor modifications.

### Transcript detection and semiquantitative RT-PCR

RNA was isolated from 100–120 mg of 14-day-old plants using the RNeasy kit (Qiagen). For cDNA synthesis, 2.5 μg of isolated mRNA was used. For PCR amplification, 2.5 μL of 20 × diluted cDNA (2 μg/ μl) was used, and the gene-specific pairs of primers were used for semiquantitative PCR (Supplementary Table S1).

### Microscopy

Microscopic observation of Arabidopsis plantlets attacked with pathogens was performed using an inverted spinning disk confocal microscope with a high-resolution camera (Yokogawa CSU-X1 on Nikon Ti-E platform, laser box Agilent MLC400, with sCMOS camera Andor Zyla CSU-X1). The dynamic study was performed with a high-speed camera (with sCMOS Andor iXon DU-897) using filter cubes for GFP. Nikon Plan Apochromat x60 WI (NA = 1.2) and Plan Apochromat x100 OI (NA = 1.45) objective lenses were used for imaging with 488 and 561 nm laser lines. The exposure time was 700 ms, with a 488 nm laser power of 75%. Fluorescence profiles and acquired images were exported from NIS ELEMENTS 4.1 software (Nikon, Tokyo, Japan); identical settings were used for each image. Images were then analysed using ImageJ Fiji software (http://rsbweb.nih.gov/ij/) and assembled with the freeware Inkscape programme into figures. Microscopic analysis of tissue staining with aniline or trypan blue was performed using an Olympus BX51 microscope with an attached DP50 camera x100 OI (NA = 1.35) objective (Olympus; Trypan blue) or Zeiss AxioImager ApoTome2 microscope 20x objective (Aniline blue). For the analysis of callotic spots, the tile images obtained with the Zeiss AxioImager ApoTome2 microscope under a 20x objective were analysed using Fiji software using the parameters of circularity of 0.5 and particle size from 200-1500 px. To exclude spots from non-specific precipitates of aniline blue in dead cells from the analysis, the cell state was assessed under bright field prior to quantification. Images obtained using the Olympus camera were analysed with analysis software and those obtained using Zeiss microscopes were analysed with ZEISS ZEN BLUE software. To perform lambda scans, a Zeiss LSM 880 confocal scanning microscope with a Zeiss C-Apochromat 40x (NA = 1.2) W Korr FCS M27 or C-Apochromat x63 OI (NA = 1.45) objective was used. For visualising the fungal structure in vivo, propidium iodide PI 1:500 or FM4-64 dye 1:1000 diluted in water was used. The excitation wavelengths were 488 nm for GFP and 561 nm for mRuby2, PI and FM4-64. Linear unmixing was performed with ZEISS ZEN BLACK software.

### Yeast two-hybrid assay

The SYP121 DNA (obtained from Riken) was amplified from cDNA with primers excluding the transmembrane domain and cloned into the pGADT7 vector (Clontech Laboratories, Inc.). All other exocyst constructs used in the study have been previously described [38,48,73](Hála *et al.*, 2008; Pečenková *et al.*, 2011; Vukašinović *et al.*, 2014). Different pGBKTs with pGAD with an inserted non-coding piece of the pENTR3C vector were used as negative controls. At least 10 positive colonies from −Leu/−Trp plates were resuspended in 150 μl of sterile water, diluted 30x and 900x, and subsequently plated on −Leu/−Trp/−His/−Ade plates.

### Protein extraction, SDS gel electrophoresis and Western blotting

Total protein extracts were isolated from 2–4-week-old plants transformed with EXO70B2-GFP at different time points ranging from 0–24 hpi with Bg or water (mock). The protein extraction Sec6/8 buffer adjusted for exocyst extraction was used (20 mM HEPES, pH 6.8, 150 mM NaCl, 1 mM EDTA, 1 mM DTT, and 0.5% Tween, supplemented with protease inhibitor cocktail (Sigma-Aldrich)). After one hour of lysis, the extracts were spun down, and the supernatants were boiled in 6x SDS loading buffer. The proteins were loaded for 10% SDS-PAGE and blotted onto a nitrocellulose membrane, with 20 ng per lane. The membrane was stained with Ponceau S solution and blocked overnight with 5% nonfat dry milk in PBS (137 mM NaCl, 2.7 mM KCl, 10 mM Na2HPO4, and 2 mM KH2PO4, pH 7.4, 0.25% Tween 20). Primary antibody dilutions in PBS were as follows: polyclonal rabbit anti-GFP (Invitrogen), 1:800; polyclonal rabbit antibodies anti-AtSEC3, 1:5000; anti-AtSEC5, 1:5000; anti-AtSEC6 (Agrisera Sweden), 1:10000; anti-AtSEC8 (Agrisera Sweden), 1:8000; anti-AtSEC10, 1:10000; anti-AtSEC15a, 1:1000 and anti-AtEXO84b, 1:1000. Appropriate secondary horseradish peroxidase conjugated antibodies (Promega, Madison, WI, USA) were applied, followed by chemiluminescent ECL detection (Amersham, GE Healthcare, Chicago, IL, USA).

### Co-immunoprecipitation

Arabidopsis seedlings (1 g of 10-day-old seedlings) were used for the co-immunoprecipitation of proteins. Protein complexes with GFP-SYP121-GFP, EXO70B2-GFP, GFP-EXO70B2 and free-GFP as a control were isolated using the μMACS GFP-tagged protein isolation kit (Miltenyi Biotec) according to the manufacturer’s instructions. The only exception was the utilisation of Sec6/8 buffer (Hála *et al.*, 2008), for experiments with GFP-SYP121 and free-GFP, as a wash buffer two times and the lysis buffer provided in the kit as a third more stringent wash (150 mM NaCl, 1% Triton X-100, 0.5% sodium deoxycholate, 0.1% SDS, 50 mM Tris HCl (pH 8.0) and subsequently a second wash buffer (20 mM Tris HCl (pH 7.5)). The elution was performed using the elution buffer provided in the kit (50 mM Tris HCl (pH 6.8), 50 mM DTT, 1% SDS, 1 mM EDTA, 0.005% bromophenol blue, 10% glycerol). To analyse bound proteins in an eluted fraction, proteins were separated using a 10% mini SDS-PAGE gel at 160 V for 20 min. The samples intended for immunodetection were processed for western blotting and immunochemical detection. The samples intended for HPLC analysis were prepared as follows: the SDS-PAGE gel was stained with colloidal Coomassie Brilliant Blue (Imperial Protein Stain, Thermo Scientific), and the whole lane was cut into four parts and submitted to in-gel digestion. The gel bands were cut into 1-mm fragments that were distained using a mixture of acetonitrile and 0.1 M ammonium bicarbonate in a 1:1 (v/v) ratio. The distained gel fragments were dried with acetonitrile, and a dithiothreitol (DTT) solution was added to reduce disulphide bonds (at 56°C for 45 min). After removal of the DTT solution, the cubes were cooled down, and an iodoacetamide solution was added. Alkylation was performed at room temperature for 30 min in the dark. The gel fragments were then washed and finally dried in acetonitrile. A solution of trypsin (12.5 ng/μl) was added to the dried gel fragments and incubated at 4°C for 25 min. The excess trypsin solution was removed, and a fresh solution of 50 mM ammonium bicarbonate was added. Proteins were digested at 37°C for 4 h. The solution containing the peptides was then transferred to a clean tube, and sufficient 0.1% TFA was added to cover the gel fragments and stop digestion. The gel fragments were sonicated for 15 min to release any peptides still present inside the gel. After sonication, both aliquots were combined, cleaned with ZipTip C18 pipette tips, and left to evaporate.

### Mass spectrometry

Mass spectrometry measurements were conducted using a UHPLC Dionex Ultimate3000 RSLC nano (Dionex, Germany) connected to a mass spectrometer ESI-Q-TOF Maxis Impact (Bruker, Germany). Samples were dissolved in 8 μl of water/acetonitrile/formic acid (97:3:0.1%), and then 5 μl of this mixture was loaded onto a trap column Acclaim PepMap100 C18 (100 μm x 2 cm, 5 μm particle size; Dionex, Germany) with a mobile phase A (0.1% formic acid in water) and a flow rate of 5 μl/min for 5 min. The peptides were eluted from the trap column onto the analytical column Acclaim PepMap RSLC C18 (75 μm x 150 mm, 2 μm particle size) by mobile phase B (0.1% formic acid in acetonitrile) using the following gradient: 0-5 min 3% B, 5-35 min 5-35% B, 37 min 90% B, 37-50 90% B, 51 min 3% B, 51-60 min 3% B. The flow rate during the gradient separation was set to 0.3 μL/min. Peptides were eluted directly to the ESI source – Captive spray (Bruker Daltonics, Germany). All data were recorded in positive ion mode. The capillary voltage was set to 1500 V, and the dry gas temperature was 150°C, at a flow of 3 l/min. Peptides for fragmentation were selected in the range from 400-1400 m/z. Up to five precursors could be selected from one MS spectrum; dynamic exclusion was set 0.5 min. MS/MS spectra were recorded in the range from 50-2200 m/z. Peak lists were extracted from the raw data with Data Analysis version 4.1 (Bruker Daltonics, Germany), and proteins were identified using the Mascot server (version 2.4.1, Matrix Science). Mass spectrometry data were matched against the Arabidopsis thaliana database (downloaded from the UniProt website on 11 October 2016, 33705 sequences) complemented with common laboratory contaminants. The following parameters were set during the search: enzyme trypsin, allowance of one missed cleavage, tolerance of 10 ppm in MS mode, and 0.05 Da in MSMS mode. The cysteine carbamidomethylation was set as a fixed modification and methionine oxidation as a variable modification; a Mascot decoy search was used to control the false discovery rate, which was set to 1%.

## Results

### EXO70B2 is specifically upregulated in early hours after fungal attack and accumulates at attack/papillae sites

The EXO70 isoform EXO70B2 mRNA is upregulated after various elicitor treatments in Arabidopsis, and the protein is involved in the biogenesis of papillae (Pečenková *et al.*, 2011; Stegmann *et al.*, 2012). We hypothesized that the EXO70B2 isoform might be specifically employed in the anti-fungal non-host defence (Žárskỳ *et al.*, 2013). To test our hypothesis, we compared the protein expression profile of EXO70B2 with its closest paralogue EXO70B1, the developmentally most abundant isoform EXO70A1 and endogenous constitutively expressed core subunit SEC8 after Bg inoculation. For this purpose, we prepared several stable lines expressing EXO70B2-GFP under the natural promoter in the *exo70B2* mutant background, which complemented its compromised phenotype in penetration resistance (Fig. S1), as well as lines with natural promoter driven GFP-EXO70B1 [50] expression in the *exo70B1* plants complementing the *exo70B1* phenotype.

The level of EXO70B2 protein increased significantly at 4 hours post-inoculation (hpi; Fig. 1) and remained unchanged even at 24 hpi (Fig. S2). The level of EXO70A1 slightly decreased, while the level of SEC8 remained unchanged, indicative of EXO70 isoforms exchange (Fig. 1A). The GFP-EXO70B1 also responded to Bg infection by upregulation, however much less than EXO70B2-GFP. Indeed, to detect the level of GFP-EXO70B1 expression in the *exo70B1* plants, we had to triple the amount of loaded proteins (Fig. 1B) in comparison to EXO70B2-GFP (Fig. 1A). To exclude the influence of the positional effect of the GFP tag, we also analysed an N-terminal fusion GFP-EXO70B2 variant, which responded to the fungus presence similarly to EXO70B2-GFP (Fig. S2).

**Figure 1.**
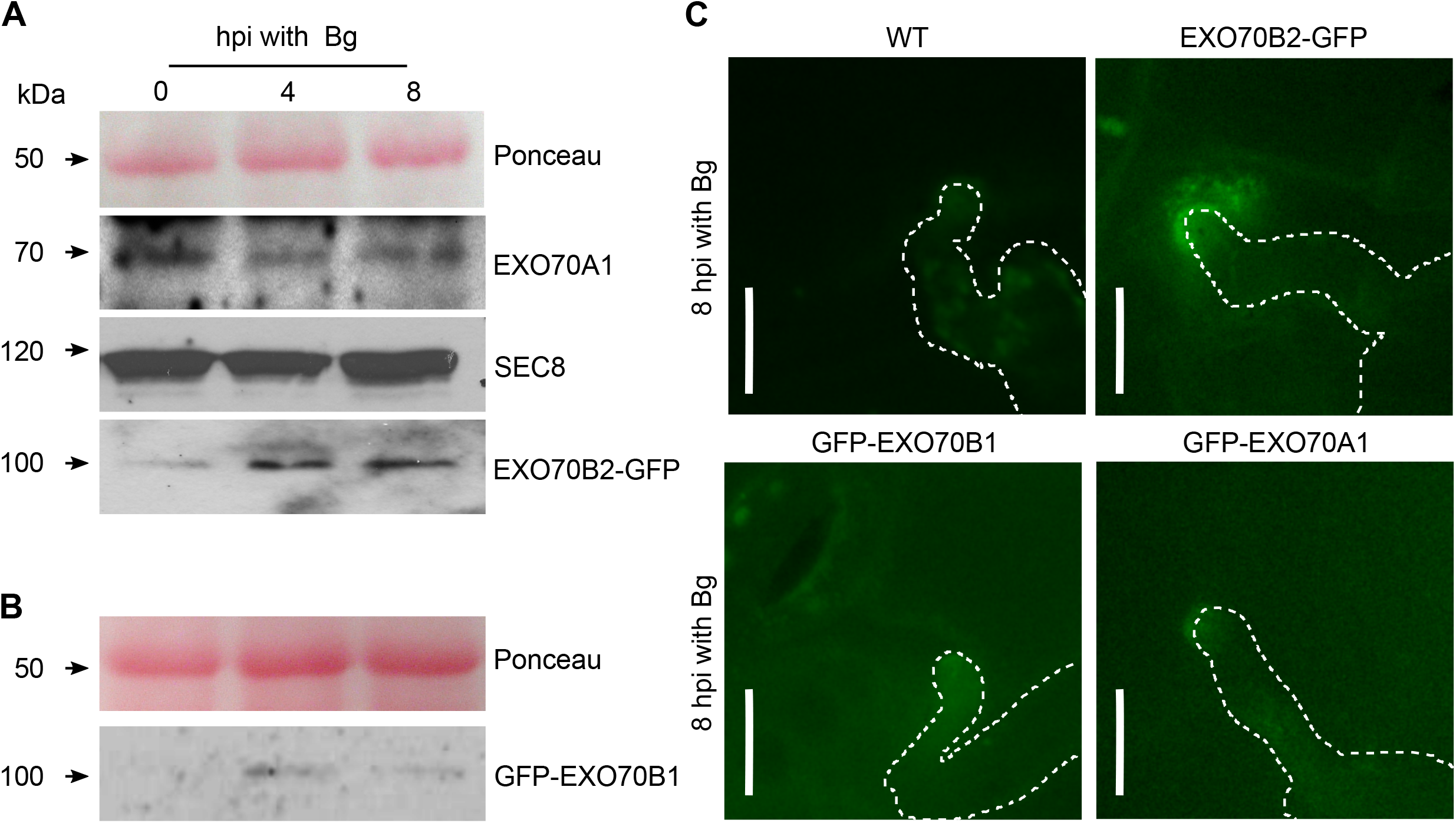
EXO70B2 expression dynamics after Bg inoculation. (A) Protein levels of EXO70A1, SEC8 and EXO70B2-GFP in the complemented *exo70B2* mutant at 0, 4, 8 hpi. (B) Protein level of GFP-EXO70B1 in the *exo70B1* mutant at 0, 4, 8 hpi. Proteins were isolated from 4-week-old plants. Anti-GFP, anti-EXO70A1 and anti-SEC8 antibodies were used. (C) The early focal accumulation of EXO70B2-GFP signal in the cytoplasm in vesicle-like dots in comparison to the absent signal of GFP-EXO70B1 and a faint signal of GFP-EXO70A1 at 8 hpi with Bg. All images represent a single optical section obtained by confocal spinning disc microscopy. All constructs were expressed under their natural promotors. The white dashed line indicates the fungus. The scale bar represents 10 μm.

To detect the earliest point of EXO70B2-GFP recruitment to the Bg/Arabidopsis contact sites, we acquired images of infected Arabidopsis leaves every two hours, starting from the 0 point of inoculation until 24 hpi. We spotted the very first signal accumulation of EXO70B2-GFP at the fungal attack sites between 8 and 9 hpi (Fig. 1C). This time corresponds to the early deposition of papillae material (Assaad *et al.*, 2004), and therefore, we propose that EXO70B2 is involved in the initial definition of the membrane domain for papilla biogenesis. In agreement with western blot analysis of protein levels (Fig. 1A), neither GFP-EXO70B1 nor GFP-EXO70A1 showed a similar early accumulation at the contact sites (Fig. 1C).

### The exocyst co-localises with the growing structure of the haustorial encasement

Despite defensive papillae-formation, non-adapted fungi such as Bg and Ep are occasionally able to successfully penetrate a non-host plant cell and develop haustoria, which later become encased, therefore we decided to analyse the presence of the exocyst subunits in both papillae structures and haustorial encasements. Using optical scanning microscopy, we inspected the surroundings of the surface of a Bg haustorium using EXO70B2-GFP and as a representative of a core exocyst subunit SEC6-GFP expressing Arabidopsis lines 48 hpi with Bg. Both tagged subunits were found to label the area of papillae (Fig. 2A, B), but also the encasements formed around Bg haustoria (Fig. 2 C, D).

**Figure 2.**
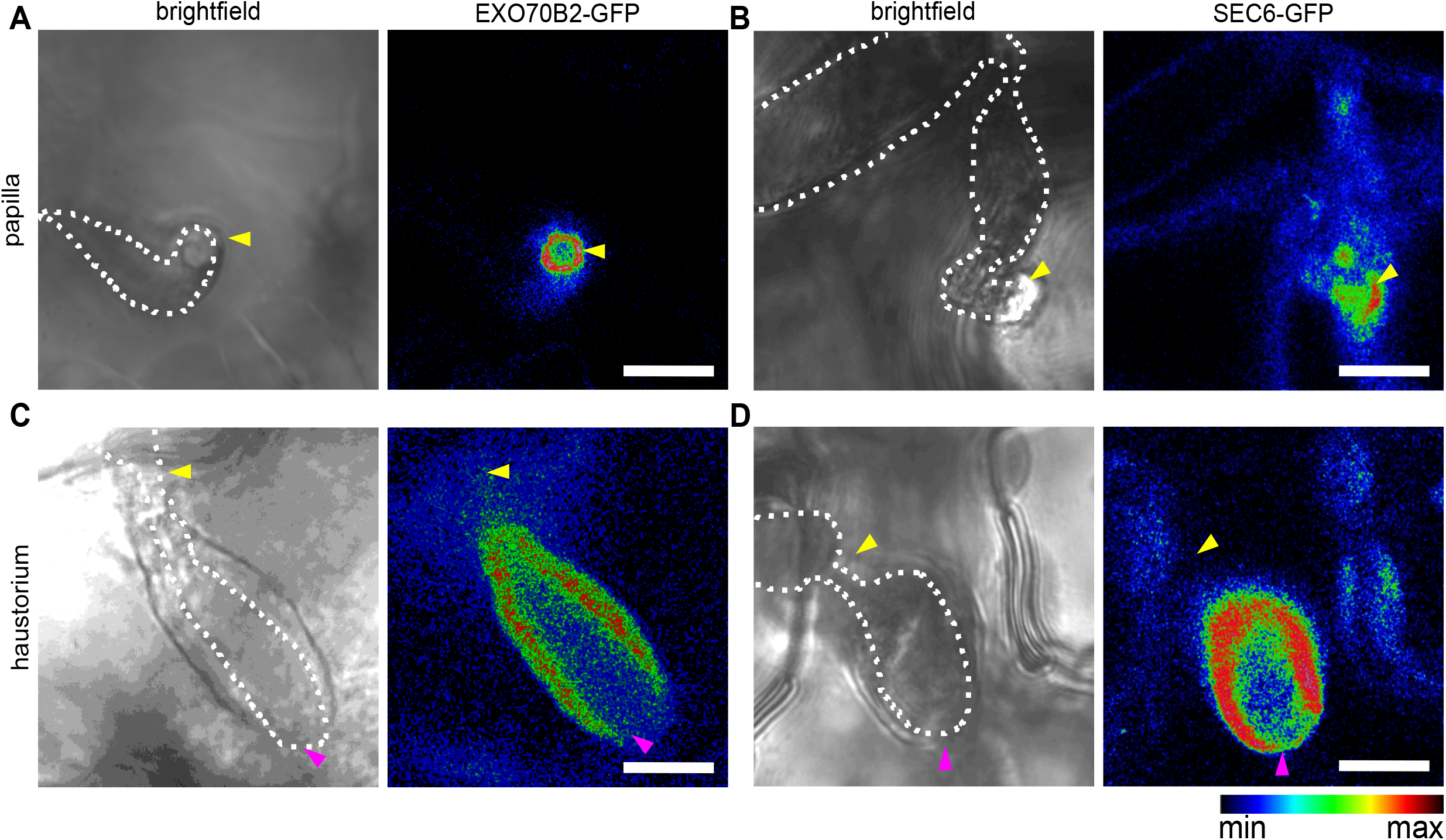
Exocyst localisation in areas of papillae and haustorial encasements. The panels represent brightfield and signal intensity analysis of the three stages of defensive structure growth: papilla (A, B) and enclosing encasement (C, D). The EXO70B2 signal surrounds the papilla (A) and the encasement (C). The SEC6-GFP signal strongly marks the papilla (B) and the encasement (D). Plants were treated for 48 hours with *Bg*. Pink arrowheads denote the haustorium encasement ends, where the exocyst subunit signal disappears. The white dashed line outlines fungal structures. The yellow arrowheads indicate penetration sites. All images represent a single optical section obtained by confocal microscopy. The scale bar represents 10 μm; the scale bar of signal intensities on the right.

The EXO70B2-GFP signal outlined the papillary body (Fig. 2A), the encasement (Fig. 2C), but was excluded from the part of haustoria lacking the encasement, i.e., it was not present at the extrahaustorial membrane (Fig. 2C, pink arrowheads). SEC6-GFP displayed a generally more intense signal than EXO70B2-GFP, in case of both papillae and haustoria (Fig. 2B and D). The exocyst subunit localisation in papillae and haustorial encasements was further verified by lambda scanning and subsequent linear unmixing analysis (Fig. S3). These observations suggest that the EXO70B2-containing exocyst complex participates in both papilla and encasement biogenesis. We also inspected the dynamics of the EXO70B2-GFP and SEC6-GFP signals within the papillae 24 hpi with Bg. The kymographs generated from 30 s of 2-min-long time series showed that the localisation of both EXO70B2-GFP and SEC6-GFP exocyst subunits within a papilla is stationary (Fig. S4), similarly to previously observed localisation of Qa-SNARE SYP121 (Nielsen *et al.*, 2012).

### Exocyst subunit mutants display abnormal papillae and impaired penetration resistance

To further investigate exocyst role in non-host resistance, we decided to analyse the defence reaction of LOF exocyst mutant plants. We employed mutants that lack strong developmental defects, such as the knockout line (KO) *sec5a-1* (GABI_731C01, Fig. S5) and the truncated mutant line *sec8-m4* (Cole *et al.*, 2005). We compared their performance with the *exo70B2* KO line and as the control, we used Col-0 ecotype. By staining with trypan blue and aniline blue, we detected, along with the previously characterised and reported types of fungal propagation (regular papilla, encased haustorium and unencased haustorium; (Takemoto *et al.*, 2006)) additional papilla structures with a vesicular halo as described previously (Pečenková *et al.*, 2011), as well as enlarged papillae with a diameter more than twice that of a regular papilla (Fig. 3, S6).

**Figure 3.**
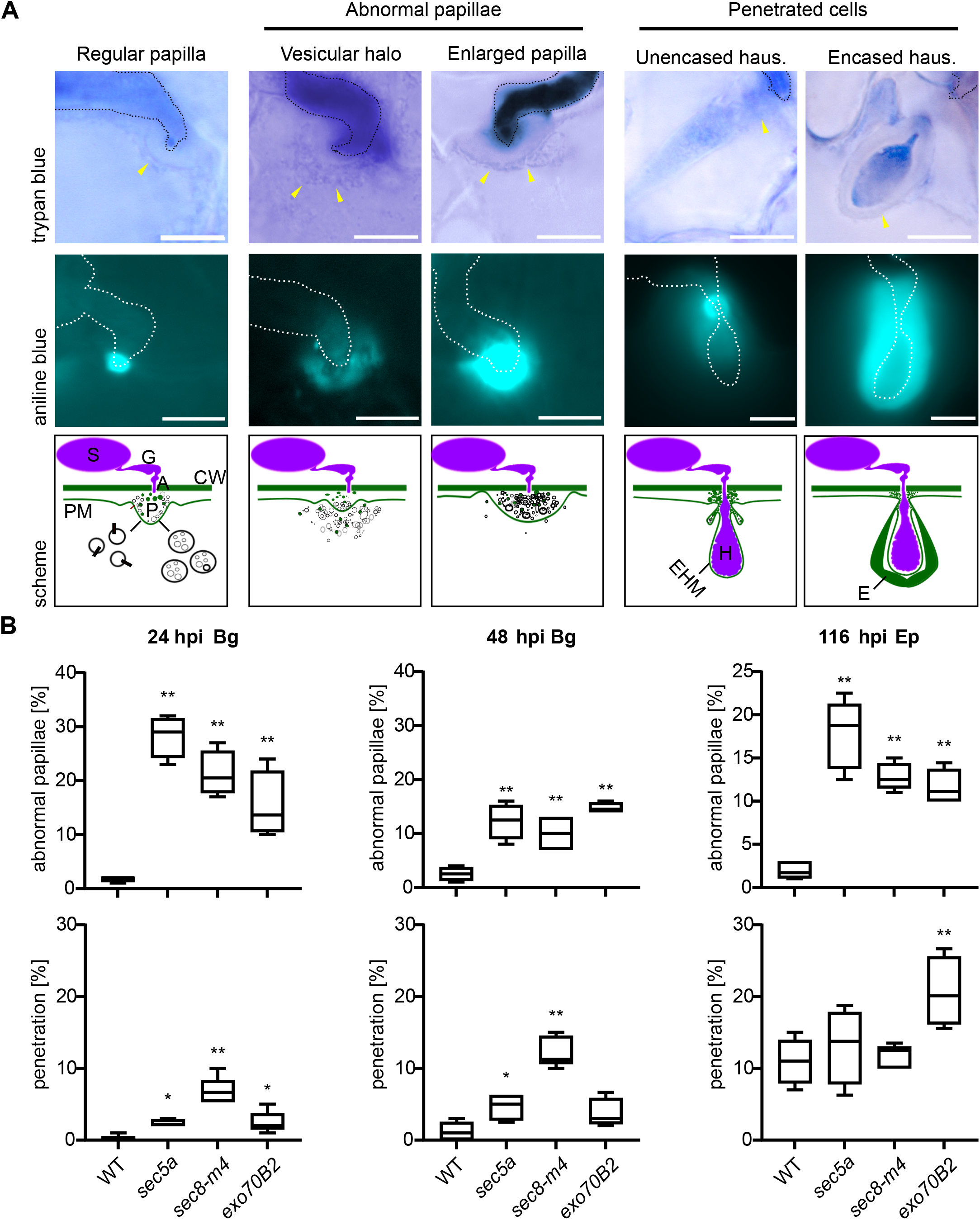
Category and quantification of the interactions between exocyst mutants and non-adapted pathogens. From the top: the first row shows independent Bg (Ep)/Arabidopsis interactions visualized by trypan blue staining; the second row displays aniline blue staining; the third row represents a schematic model of the interaction. Bg and Ep evoke the same types of interactions in Arabidopsis: regular papilla; abnormal papillae / either with a vesicular halo or enlarged papilla; penetrated cells / either unencased or encased haustorium. (B) The data show the mean of the abnormal papillae and penetrated cells by tested pathogens at different time points. For each column in the graph, the area complementary to 100% represents all interactions with spores that did not provoke any reaction of the plant. The experiment was repeated 3 times with a similar trend. In each experiment, the N represents 5 quantified leaves, 100 spores were counted per leaf. Stars indicate significant differences calculated by one-way ANOVA with the post-hoc Tukey-Kramer Honestly Significant Difference (HSD) test, ** Z-score p<0.01, * Z-score p<0.05. For the schematic: A appressorium; GT germ tube; S spore; CW cell wall; PM plasma membrane; P papilla; EHM extrahaustorial membrane; H haustorium; E encasement. Yellow arrows mark outer borders of plant cell defensive structures. The dashed line outlines fungal structures. Scale bars represent 5 μm.

In these abnormal papillae, we identified two distinct patterns of callose deposition. The vesicular clump category exhibited a reduction of callose and the stacking of faint callose spots around the appressorium (Fig. 3A, S7). In contrast, the enlarged papillae showed an over-accumulation of callose, as well as a larger papilla body observed under the bright field (Fig. 3A, S7). We quantified the occurrence of these papillae and penetration efficiency at 24 and 48 hpi with Bg and 116 hpi with Ep in the selected mutant lines (due to the slower spore germination of Ep, we prolonged the inoculation time for evaluation of its penetration; Fig. 3B). High penetration efficiency was correlated with cells either developing haustoria or undergoing cell death (Fig. 3A). In accordance with published data (Pečenková *et al.*, 2011), *exo70B2* plants demonstrated a higher occurrence of deviated papillae, as well as an increased penetration efficiency of Ep (Fig. 3B). The *sec5a-1* and *sec8-m4* mutant lines showed significant increases in both the deviated papillae and the penetration rate (Fig. 3B).

### Time course comparison of callose deposition and haustorium formation between exocyst and syp121 mutants

The observed morphological changes of papillae indicate EXO70B2-containing exocyst complex (including SEC8) is involved in papillae biogenesis and callose deposition within papillae. In order to quantify a range of the exocyst disruption effect on the callose deposition, we quantified the Bg-induces callose punctae on leaves of mutants *exo70B2* and *sec8-m4* in comparison to the wild type control, and also LOF *syp121*, a mutant of the SNARE protein that has been shown to be important for timely deposition of callose into papillae (Collins *et al.*, 2003; Assaad *et al.*, 2004) (Fig. 4A). In addition, we followed the time course development of haustoria in living cells in these lines (Fig. 4B).

**Figure 4.**
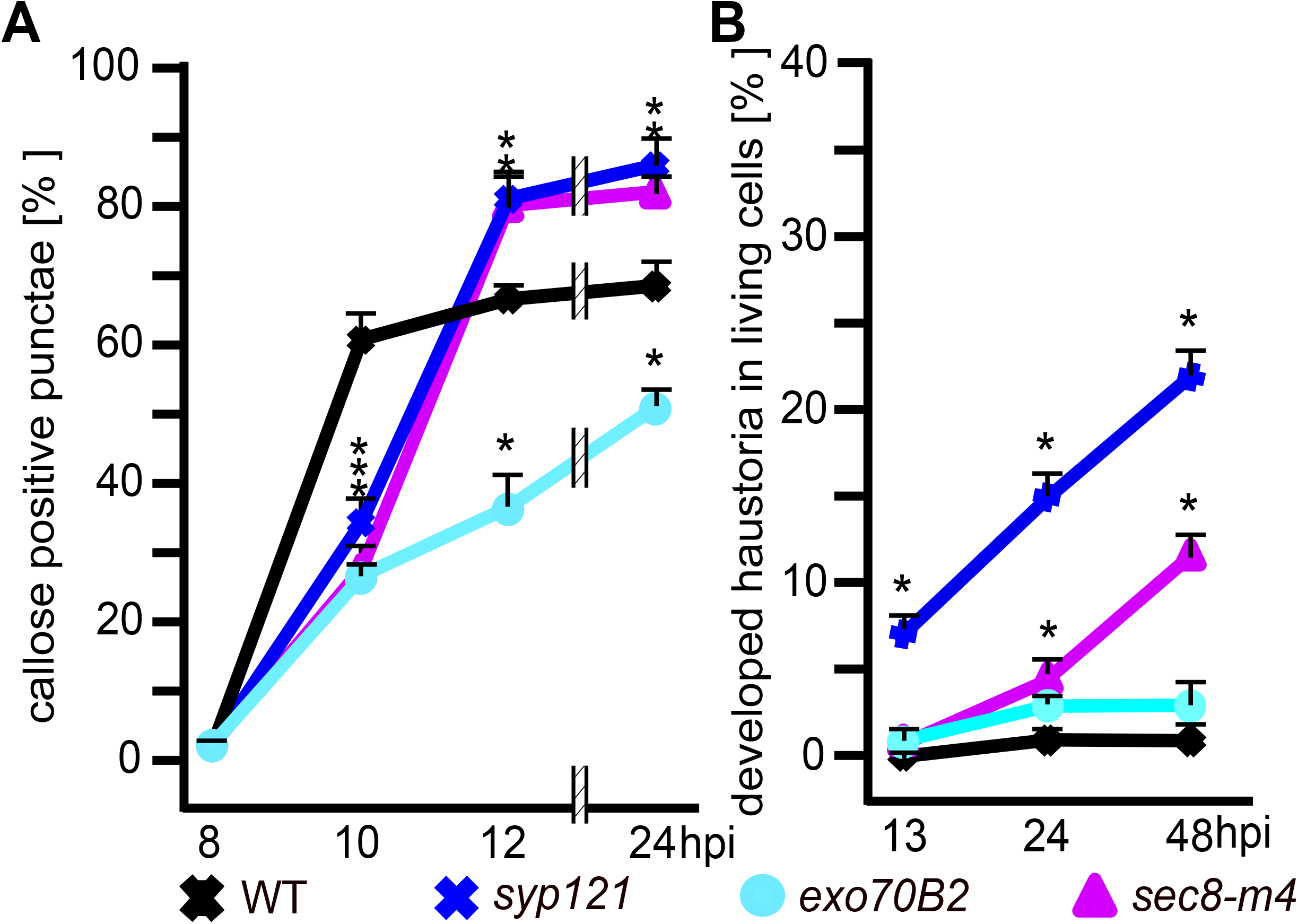
Effects of exocyst mutations on the occurrence of a haustorium and callose deposition in defence against Bg. (A) The frequency of callose positive punctae counted on a defined leaf area at sites of Bg attack was followed at 8, 10, 12, 24 hpi. (B) The frequency of developed haustoria in cells of WT, *syp121*, *exo70B2* and *sec8-m4* leaves. Each experiment was repeated 3 times with similar results, N represents 5 leaves. The asterisks indicate significant difference from WT at each time point. Statistical significance was calculated using one-way ANOVA with the post-hoc Tukey-Kramer HSD test, Z score p<0.05. Error bars represent the standard error.

In our experiment, the *sec8-m4* mutant displayed faster penetration by Bg than the WT, but still slower penetration than the *syp121* mutant (manifested as formed haustoria number; Fig. 4B). The higher penetration success of Bg in *syp121* has been associated with the delay in callose deposition in early papilla development. We observed a comparable delay in callose deposition for all three *sec8-m4*, *exo70B2* and *syp121* single LOF mutants at 10 hpi. Surprisingly, the *exo70B2* mutant exhibited the strongest decrease in callose deposition, which remained apparent until the later time points after inoculation.

### EXO70B2 can interact with SYP121

Results of the analysis of callose deposition indicate a functional overlap and a connection between the EXO70B2/exocyst- and SYP121-driven secretory pathways, similarly to the situation in yeast, and also in seedlings of *A. thaliana* (Sivaram *et al.*, 2005; Larson *et al.*, 2020). To examine the possibility that the two complexes might interact in the non-host anti-fungal defence via EXO70B2-SYP121 direct interaction, we employed a yeast-two hybrid system. We used the truncated construct SYP121ΔC (free of the transmembrane domain) and besides EXO70B2 also several other exocyst complex subunits (AD-SYP121ΔC/BD-EXO70A1, EXO70B1, EXO70B2, EXO70H1, SEC3a, SEC5a, SEC6, SEC8, SEC10a, SEC15b, EXO84b, and the pair SEC3a/EXO70A1 was used as a positive control; Fig. 5A). We detected only a weak interaction of SYP121ΔC with EXO70B2 and EXO70B1 (A). To verify the interaction in planta, we examined the GFP-SYP121 bound fraction with available anti-exocyst antibodies. In the eluate, we detected the presence of two exocyst core subunits, SEC6 and SEC3 (Fig. 5B), confirming the ability of the exocyst complex to associate with SYP121 in Arabidopsis. However, using the opposite approach, where EXO70B2-GFP after plant treatment with Bg was used as bait for co-immunoprecipitation followed by sensitive HPLC MS/MS analysis, did not reveal the presence of SYP121 (Supplementary Table 2).

**Figure 5.**
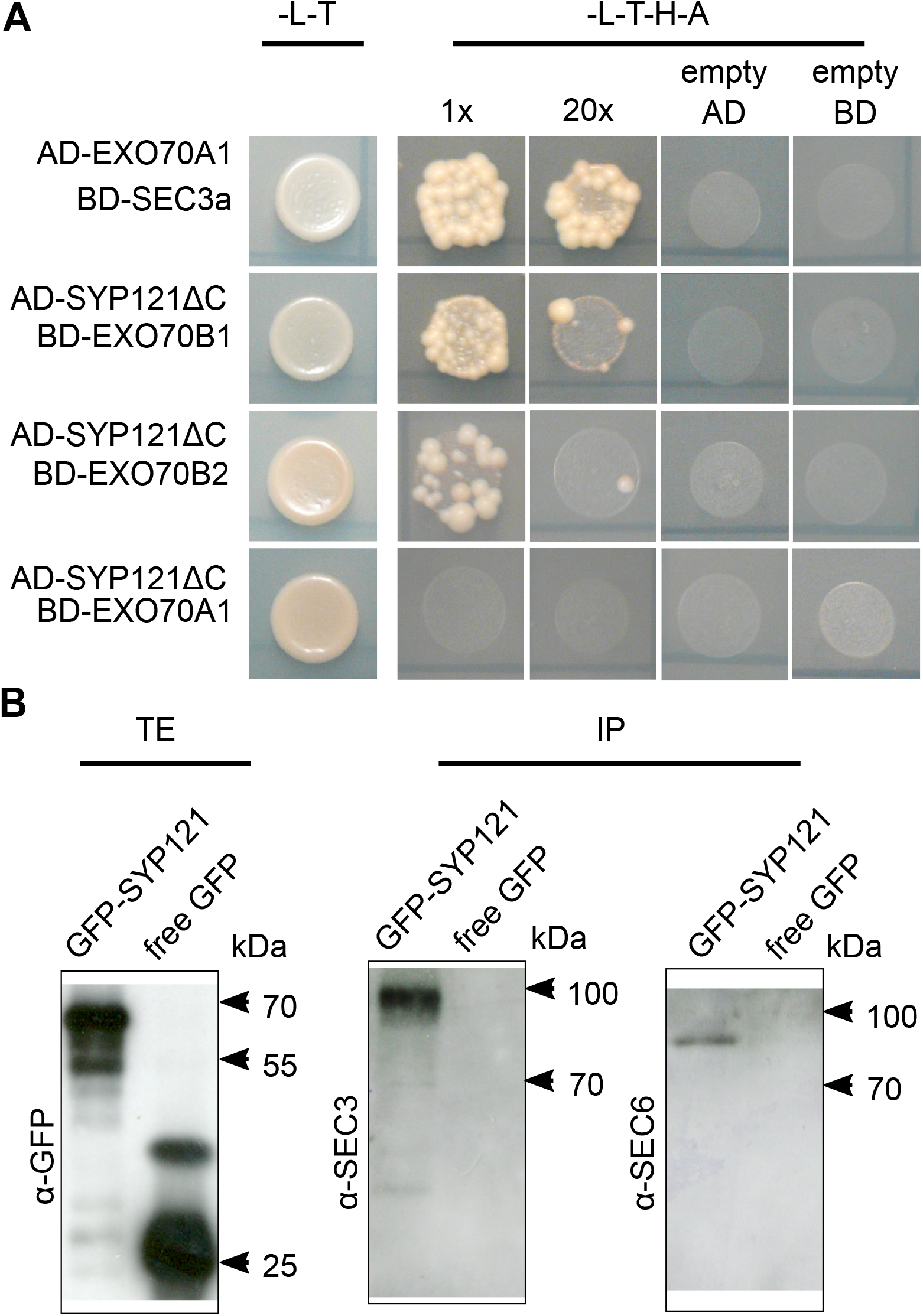
EXO70B2 subunit interacts with SYP121 in defence. (A) Direct interaction between the exocyst and SYP121 was examined in a yeast two-hybrid assay. Yeasts containing both the binding (BD) and activation domain (AD) were plated on the double selection SD-Leu, -Trp or on the SD-His, Trp, Ade, Leu plates and allowed to grow for 6 days. For each pair, negative control transformations with empty counterpart AD or BD plasmids were performed. (B) The seedlings expressing GFP-SYP121 or free GFP were used for anti-GFP co-immunoprecipitation and the bound fraction was tested for the exocyst subunits presence by immunoblotting. TE; total extract, IP immuno-precipitated proteins.

### EXO70B2 and SYP121 cooperate in papillae biogenesis

Since EXO70B2 and SYP121 both regulate early papillae biogenesis and are capable of direct interaction, we searched for evidence of their functional cooperation. Using the stable transgenic Arabidopsis lines, we examined the co-localisation of mRuby2-EXO70B2 with GFP-SYP121 16 hpi with Bg (Fig. 6A). Both proteins co-localised at the papilla periphery (white colour), and their fragile co-localisation disappeared after plasmolysis (Fig. S8). Unlike GEP-SYP121, mRuby2-EXO70B2 was absent from the apoplastic space of papillae (Fig. S8).

**Figure 6.**
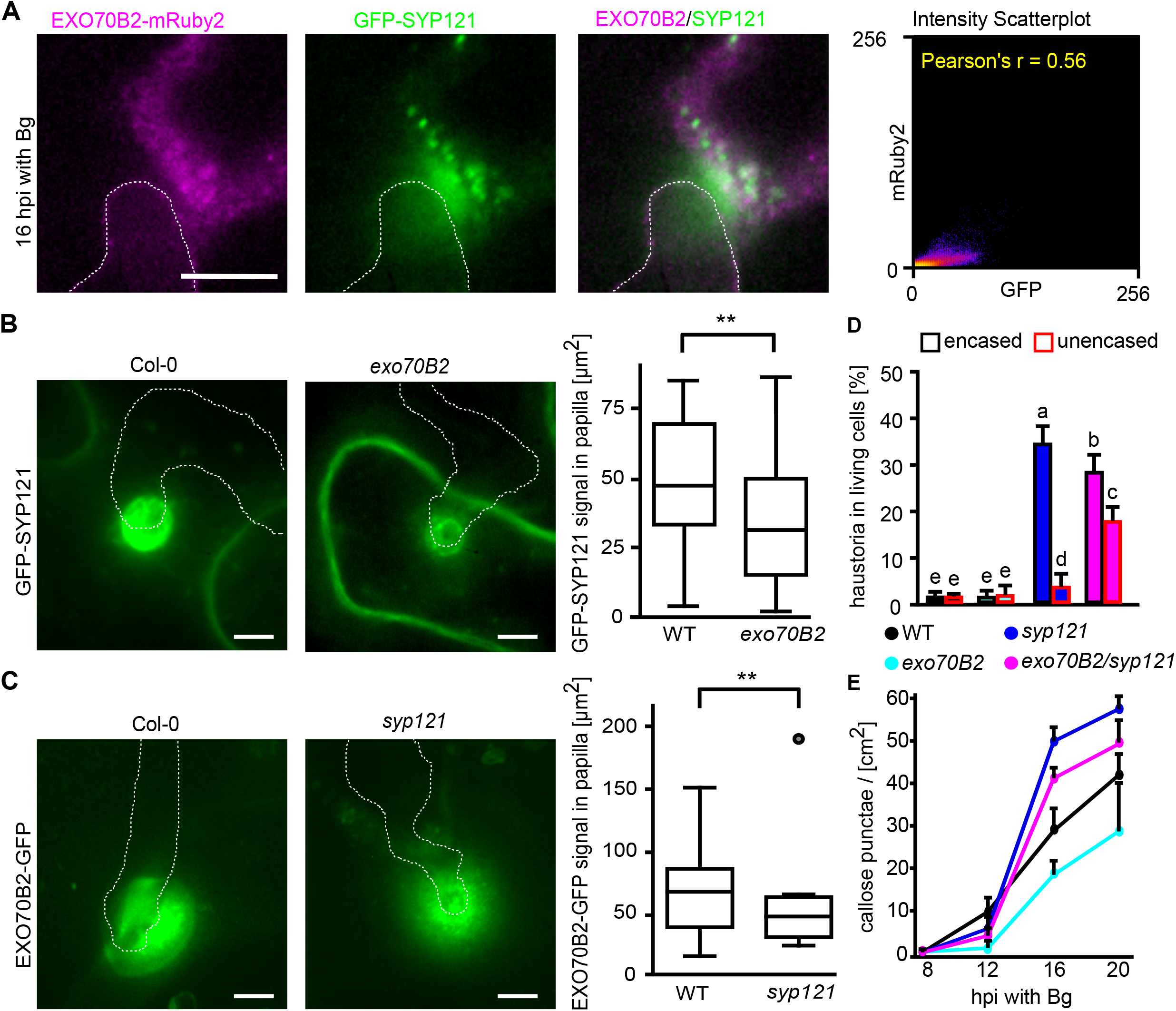
EXO70B2 subunit and SYP121 SNARE cooperate in papillae biogenesis. (A) Co-localisation of mRuby2-EXO70B2 and GFP-SYP121 in the membrane domain of a defensive papilla. The transgenic plants with the mRuby2-EXO70B2 and GFP-SYP121 were observed at 16 hpi with Bg; images represent one of 15 analysed papillae with similar localisation. Pearson’s correlation graph of mRuby2-EXO70B2 (Ch1) and GFP-SYP121 (Ch2) is shown on the right. (B) Representative images and quantification of GFP-SYP121 signal area in papillae in Col-0 and *exo70B2* mutant backgrounds. The N is equal to 30 Z-stack images, consistent with 12.5 um thickness, for each genotype. The data were processed via two-tail T-test on p<0.01 value. (C) Complementary to panel B, the image and quantification of EXO70B2-GFP signal area in Col-0 and *syp121*. Analysed with the same parameters as in B. The white dashed line marks the cross-section used for analysis. All images represent single plane sections. The scale bars represent 5μm. (D-E) Analysis of Col-0, *syp121*, *exo70B2* and *exo70B2/syp121* mutant reactions to Bg infection. Quantification of encased and unencased haustoria present in living cells, N is equal to 5 leaves, 100 spores were counted per leaf. Small letters indicate statistical differences analysed by ANOVA post-hoc Tukey-Kramer (HSD) Z score p<0.01. Error bars represent standard deviations. (E) Accumulation of callose punctae beneath fungal contact sites per square cm of leaf. The N is equal to 5-8 leaves. The error bars represent standard deviations.

We then inspected the localisation patterns of EXO70B2 and SYP121 in counterpart mutant lines backgrounds. First, we followed the localisation of GFP-SYP121 in papillae at 24 hpi with Bg in WT and *exo70B2* mutant plants (Fig. 6B). In the WT background, the maximum intensity projection of GFP-SYP121 signal appeared due to its extracellular localisation as a filled circular area of variable diameter (Fig. 6B), as described previously (Collins *et al.*, 2003; Assaad *et al.*, 2004). However, in the *exo70B2* mutant background, GFP-SYP121 labelled a significantly smaller papilla area (Fig. 6B). We also noticed that the GFP-SYP121 signal in *exo70B2* plants often showed a thin doughnut-like pattern (hollow circle) at plant/fungal contact sites, indicative of missing extracellular localisation as observed in the WT (Fig. 6B). Second, we analysed the localisation of EXO70B2-GFP in the WT and *syp121* mutant background. In this case, we did not observe a change in the pattern of the signal itself, but the area of the EXO70B2-GFP papillae was smaller in *syp121* compared with WT (Fig. 6C). We decided to further test the relevance of the interaction of EXO70B2 and SYP121 to penetration resistance at the genetic level. We examined the penetration resistance of the double mutant *exo70B2/syp121* to Bg in comparison to the single *syp121* and *exo70B2* mutants (the appearance of the infected plants is shown in Fig. S9). The *exo70B2/syp121* double mutant had a significantly decreased number of regular papillae in comparison to the single *syp121* (Fig. S9). Despite an absence of a significant difference in the total number of haustoria in living cells (Fig. S9), the *exo70B2/syp121* double mutant had a decreased number of encased haustoria and an increased number of unencased haustoria compared with the *syp121* single mutant (Fig. 6D). In our set of tested mutants, we observed that both single and double mutants showed similar delays in callose deposition at earlier time points p.i.; however, at later time points the additional loss of EXO70B2 in *syp121* slightly but steadily dampened the otherwise increased level of callose punctae (Fig. 6E).

## Discussion

### Exocyst is present in membrane domain of papillae and encasements around haustoria

The exocyst roles in regulation of plant cell secretion and polarity, and also surprising involvement in autophagy, imply its participation also in defence responses (Žárskỳ *et al.*, 2013). Indeed, in accord with the published observations of EXO70B2, the isoform of EXO70 subunit of the exocyst complex, mRNA upregulation by PAMPs (Pečenková *et al.*, 2011; Stegmann *et al.*, 2012), we documented early protein upregulation and focal accumulation of EXO70B2-GFP and core exocyst subunit SEC6-GFP in areas of Blumeria penetration attempts and papilla and haustorium encasement formation (Fig. 1, 2). We showed that this enhanced localisation was a specific regulated response since the increase in the EXO70B2-GFP protein level correlated with mild EXO70A1 depletion, while the level of another core subunit SEC8 remained unchanged (Fig. 1A). Our data also support the specific role of EXO70B2 in the biogenesis of papillae over its closest paralogue, the EXO70B1, while, as has been shown recently, both cooperate in FLS2 signalling (Wang *et al.*, 2020). Clearly, the localisation and expression pattern shows differences from the rest of the complex, which, along with the previous results documenting its ability to interact with exocyst core, suggests that EXO70B2 employ only proportionate part of the core subunits. Interestingly, for the analysed exocyst subunits, the localisation in the membrane domain of papillae was found to be persistent over time (Fig. S4). A similarly stable localisation of exocyst subunits has been observed during secondary cell wall deposition in tracheary elements or trichomes (Kulich *et al.*, 2015, 2018; Vukašinović *et al.*, 2017), in contrast to the dynamic exocyst localisation at the lateral PM domains of rhizodermal cells (Fendrych *et al.*, 2013). These data suggest that exocyst tends to stay trapped in secretory domains involved in deposition of cell-wall components, particularly callose which is a component of both papilla and encasement around a haustorium.

### Impaired balance of the exocyst complex compromises penetration resistance

A disruption of the plant exocyst complex leads to severe growth phenotypes (Synek *et al.*, 2006; Hála *et al.*, 2008; Kulich *et al.*, 2013; Žárskỳ *et al.*, 2013). To study the specific role of the exocyst in anti-fungal defence, but to avoid pleiotropic mutation effects of full exocyst LOF, we worked, along with the immunity-related mutant *exo70B2*, also with weak allele lines of Arabidopsis core exocyst subunits *sec8-m4* and *sec5a-1*. Using these mutants, we were able to describe new Arabidopsis-Bg interaction phenotypic deviation, apart from the previously described fungal stages found for WT (Takemoto *et al.*, 2006). In addition to the previously characterised vesiculated papillae (Pečenková *et al.*, 2011), we newly described the presence of enlarged papillae. These abnormal papillae in the exocyst mutants manifested uneven deposition of callose consisting of either faint callose signals or excessive callose deposition on the contrary (Fig. 3). Such enlarged papillae with excess callose resembled the phenotype of the Arabidopsis line overexpressing PMR4 callose synthase (Blümke *et al.*, 2013). In the core subunit *sec8-m4* mutant, we observed stronger penetration defects than in the *exo70B2* mutant. This finding may reflect a general role of SEC8 in the maintenance of the secretory efficiency, in contrast to the specific requirement for EXO70B2 in defence. Intriguingly, even though the *exo70B2* and *sec8-m4* mutants, similarly to the *syp121* mutant, exhibited a delay in early callose deposition, their penetration resistance was not compromised as drastically as in the case of *syp121* (Fig. 4). We could speculate that these milder phenotypes in penetration resistance were the result of a compensation by another EXO70 isoform, as well as the partial function of the truncated SEC8 in the case of *sec8-m4*. On the other hand, there are multiple reports on Arabidopsis mutants defective in callose deposition whose overall resistance is not compromised (Aist, 1976; Jacobs *et al.*, 2003; Nishimura *et al.*, 2003), therefore our data might also support a view that the deposition of callose may not be of key importance for the final defence outcome. This assumption is supported by another observation, wherein the double mutant *exo70B2/syp121* was the deposition of stress-related callose dampened even more than in *syp121* single mutant, however without a major change in total numbers of penetrated cells (Fig. 6, S9).

It has been previously suggested that the callose is loaded into papillae via multivesicular bodies rather than being synthesized within the cell wall apposition (Böhlenius *et al.*, 2010). This phenomenon would imply an involvement of the exocyst in non-canonical secretion to the papillae extracellular space that would involve callose but also GFP-SYP121 as a non-functional cargo (Nielsen *et al.*, 2012), and very probably other defence-related cell wall material and phytoalexins. This theory is supported by the association of GFP tagged EXO70B2 with MVB-related RabG3C in pathogen-treated plants (Supplementary Table S2), however this needs to be verified in future, as well as if and how the EXO70B2 papillae function is related to phosphorylation status and interaction with ATG8 (Brillada *et al.*, 2021). In agreement with these observations is the employment of EXO70B2 in trafficking seemingly other than canonical exocytosis.

### The exocyst complex affects SYP121 SNARE in antifungal defence

In this report, we showed that SYP121 and EXO70B2 are capable of direct interaction, they co-localise, affect each other’s localisation and have additive defence effects in functional genetic analyses. The cooperation of the two proteins could also be extrapolated based on their co-expression and interactions with SNAP33 (Pečenková *et al.*, 2011; Bozkurt *et al.*, 2015). Interestingly, our y2h test revealed that the soluble part of SYP121 was able to interact with EXO70B1 but not EXO70A1 (Fig. 5A). Although we did not find a relevant role for EXO70B1 or for EXO70A1 in non-host resistance, we admit that other EXO70 isoforms might cooperate with SYP121 in defence. Our EXO70B2/SYP121 cooperation model, despite several positive tests, was not supported by HPLC-MS/MS analysis of proteins bound to EXO70B2-GFP. A reason for this is most probably a transient nature of the exocyst (peripheral membrane complex)–SNARE (integral membrane proteins) interaction. Regarding the relationship of EXO70B2 and SYP121, the most relevant distinction between the two proteins lies within the observation that unlike GFP-SYP121, EXO70B2-GFP is not secreted into the extracellular papillar space while it does contribute to GFP-SYP121 unconventional extracellular papillar localisation, the relevance of which remains to be analysed. Based on the interaction capability, mutual dependent localization as well as additive effect of the *exo70B2/syp121* double mutant, we conclude that EXO70B2 and SYP121 cooperate in the secretory pathway that drives cell wall and defence material towards the papillae and encasements in response to the fungal attack (Fig. 7).

**Figure 7.**
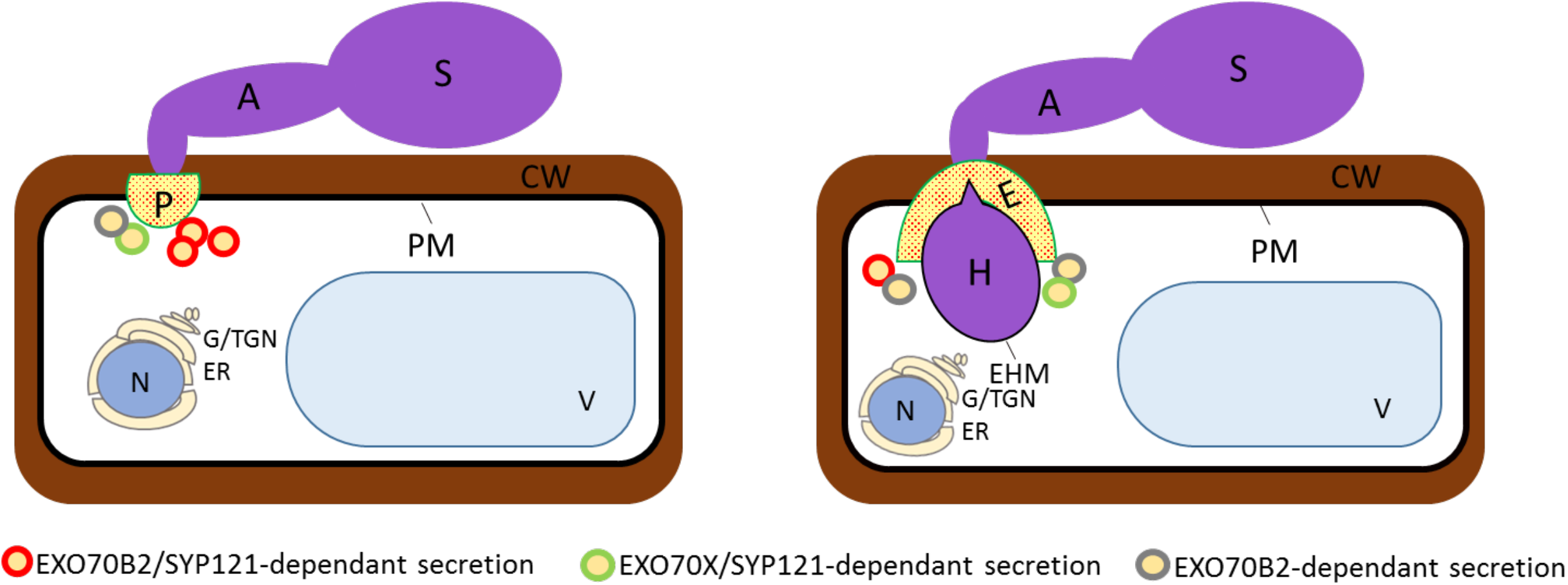
Schematic of exocyst/SYP121 cooperation in defence against Bg. The schematic on the left shows a germinated spore and host cell papilla formation; the schematic on the right shows a spore forming haustoria after successful penetration into the host cell; the host cell responds by encasement formation. The cell uses at least three different secretory pathways with respect to EXO70B2 and SYP121 involvement: EXO70B2/SYP121-dependent, EXO70X/SYP121-dependent and EXO70B2-dependent and SYP121-independent. These pathways may also include the recycling and transcytosis of vesicles. We suggest that the presence of a particular EXO70 in the complex helps the exocyst to distinguish the membrane domain of defensive structures from other PM domains. The EXO70B2-containing exocyst may help to establish the papilla membrane domain for SYP121, and similarly in encasement formation also for another SNARE participant. A appressorium, CW cell wall, E encasement, EHM extrahaustorial membrane, ER endoplasmic reticulum, G/TGN Golgi/trans Golgi network, H haustorium, N nucleus, P papilla, PM plasma membrane, S spore, V vacuole.

Our data show that the exocyst, mainly its EXO70B2-containing variant, plays a key role in papilla and haustorial encasement build-up, including callose secretion, during non-host defence. The exclusive role of the EXO70B2 subunit in papillae and encasement formation is probably executed via a functional relationship with the SYP121-carrying SNARE complex. However, in addition to the usual consideration of the existence of alternative redundant complex variants, our data do not exclude the existence of diversified exocyst and SNARE pathways involved in defence. The EXO70B2-dependent SYP121 localisation to the paramural space vs. the absence of the exocyst in it raises especially interesting questions concerning the mechanistic details and time sequence of the action of the exocyst vs. SNARE, which requires further research.

## abbreviations

Bg: Blumeria graminis f. sp. hordei
PAMP: pathogen associated molecular patterns
ROS: reactive oxygen species
EHM: extra haustorial membrane
Ep: Erysiphe pisi
PM: plasma membrane
SNARE: soluble N-ethylmaleimide-sensitive factor Attachment protein REceptors
SNAP: soluble N-ethylmaleimide-sensitive factor adaptor protein
VAMP: vesicle-associated membrane protein
MVB: multivesicular bodies
hpi: hour post inoculation
KO: knockout line
HSD: honestly significant difference
SA: salicylic acid
BD: binding domain
AD: activation domain
NA: numeric aperture

## Supplementary data

Supplementary Fig. 1. The EXO70B2-GFP and GFP-EXO70B2 complementation of the *exo70B2* mutant phenotype.

Supplementary Fig. 2. Expression level of exocyst subunits after Bg inoculation.

Supplementary Fig. 3. Linear unmixing analysis of GFP-tagged exocyst subunits in papillae and haustoria.

Supplementary Fig. 4. Exocyst dynamics in papillary membrane domains.

Supplementary Fig. 5. Verification of the KO *sec5a-1* mutant.

Supplementary Fig. 6. Detailed visualisation of Bg/Arabidopsis interactions using trypan blue staining.

Supplementary Fig. 7. Detailed visualisation of Bg/Arabidopsis interactions using aniline blue staining.

Supplementary Fig. 8. EXO70B2 does not follow SYP121 into apoplast in defensive papilla.

Supplementary Fig. 9. Analysis of *exo70B2/syp121* double mutant reaction to Bg infection.

Supplementary Table 1. List of primers and constructs used in the study.

Supplementary Table 2. Identified proteins in EXO70B2-GFP bound fraction.

## Acknowledgements

This work was supported by Czech Science Foundation (CSF/GACR) project 19-02242J and terminated GA16-34887L, a terminated student project GAUK-1102214 (Charles University Grant Agency), and the Fund-Project “Centre for Experimental Plant Biology”: No. CZ.02.1.01/0.0/0.0/16_019/0000738 of MEYS CR. The Institute of Experimental Botany (IEB) Imaging Facility is supported by the Operational Programme Prague Competitiveness (OPPC) CZ.2.16/3.1.00/21519 and Czech-BioImaging large RI project LM2015062 funded by MEYS CR.

## Author contributions

Conceived and designed the experiments: JO, TP, VZ. Performed the experiments: JO, TP. Analysed the data: JO. Contributed reagents/materials/analysis tools: JO, TP, JS, JŠ, IK, PD. Wrote the paper: JO, TP, JS, VZ

## Data availability statement

All data supporting the findings of this study are available within the paper and within its supplementary materials published online.

The data supporting the findings of this study are available from the corresponding author (Tamara Pečenková) upon request.

## Notes

### Competing Interest Statement

The authors have declared no competing interest.

