## Supplementary material for "The EXO70B2 exocyst subunit contributes to papillae and encasement formation in anti-fungal defence in Arabidopsis": Supplementary data.pdf

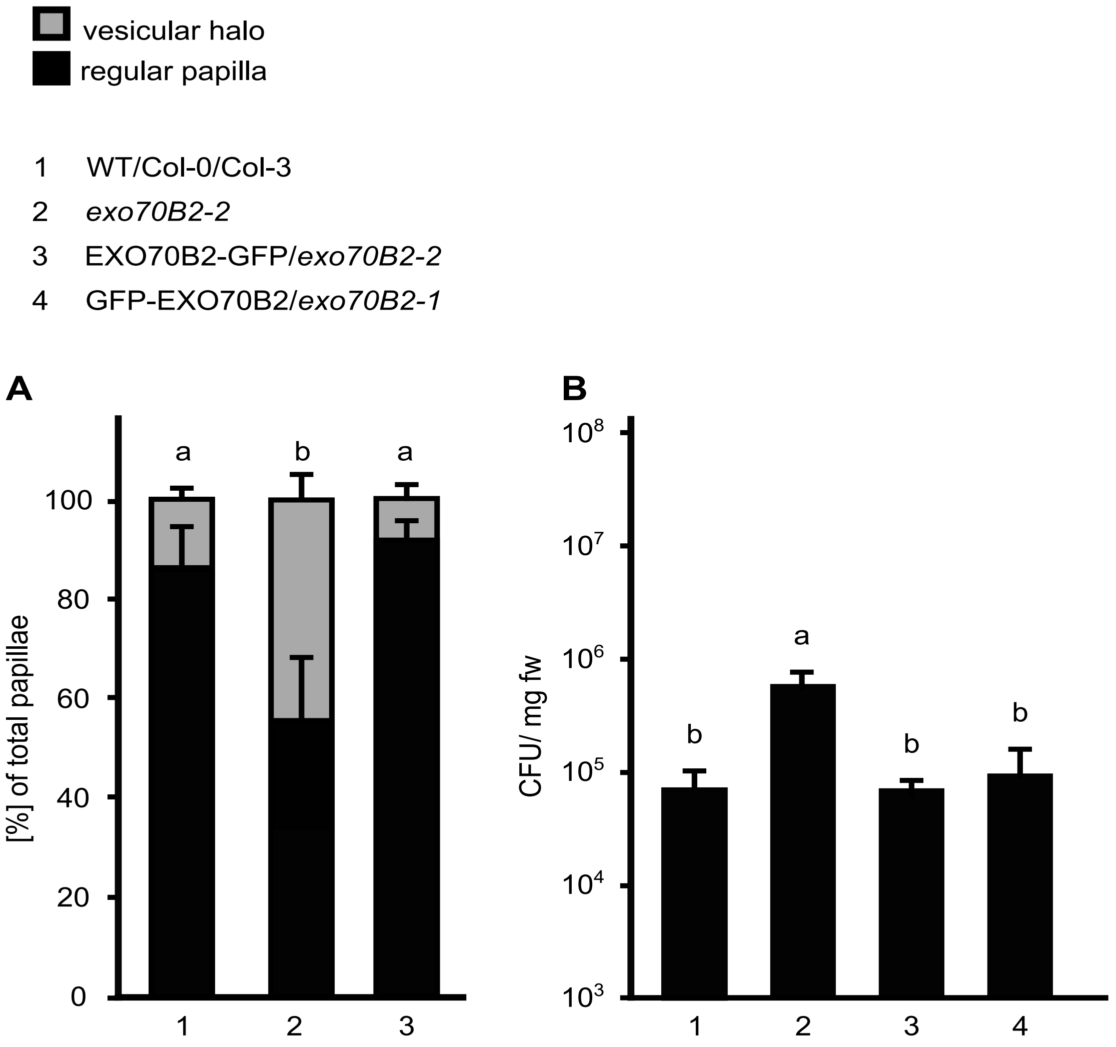


**Figure S1. The EXO70B2-GFP and GFP-EXO70B2 complementation of the *exo70B2* mutant phenotype.** (A) The EXO70B2-GFP fully complements the defect of mutant *exo70B2-2* in papilla development at 48 hpi with Bg. Leaves were stained using the trypan blue method. From the total number of created papillae on a leaf, papillae with a vesicular halo and regular papillae were counted. Error bars represent the SE. Small letters indicate statistically significant differences calculated by ANOVA followed by the Tukey-Kramer (SHD) post hoc test at $p<0.01$. n=100 spores per leaf, with 5 leaves taken from each genotype. (B) EXO70B2-GFP and GFP-EXO70B2 fully complemented the defect of the *exo70B2-2* mutant in sensitivity to *Pseudomonas syringae* pv. *tomato* DC3000 at 24 hpi in a seedling flood-inoculation assay. n=3, small letters indicate statistically significant differences calculated by ANOVA with Tukey-Kramer post hoc test at $p<0.05$.

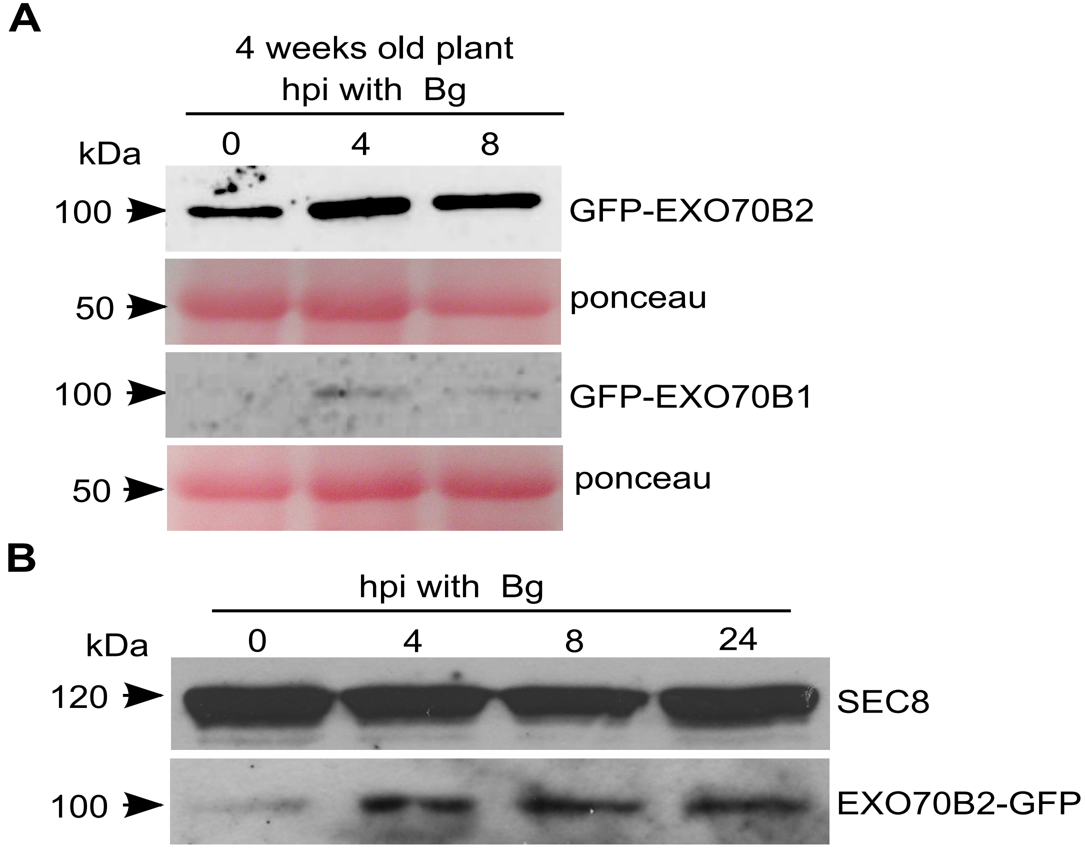


**Figure S2. Expression level of exocyst subunits after Bg inoculation.** (A) Anti-GFP antibody was used to visualize EXO70B2 and EXO70B1 protein levels at 0, 4, 8 hpi with Bg. Ponceau dye was used to visualize the amount of loaded proteins and the Rubisco large subunit as control, respectively. (B) Prolonged treatment of complemented EXO70B2-GFP plants up to 24 hpi with Bg. Anti-GFP and anti-SEC8 antibody were used to visualise exocyst subunits.

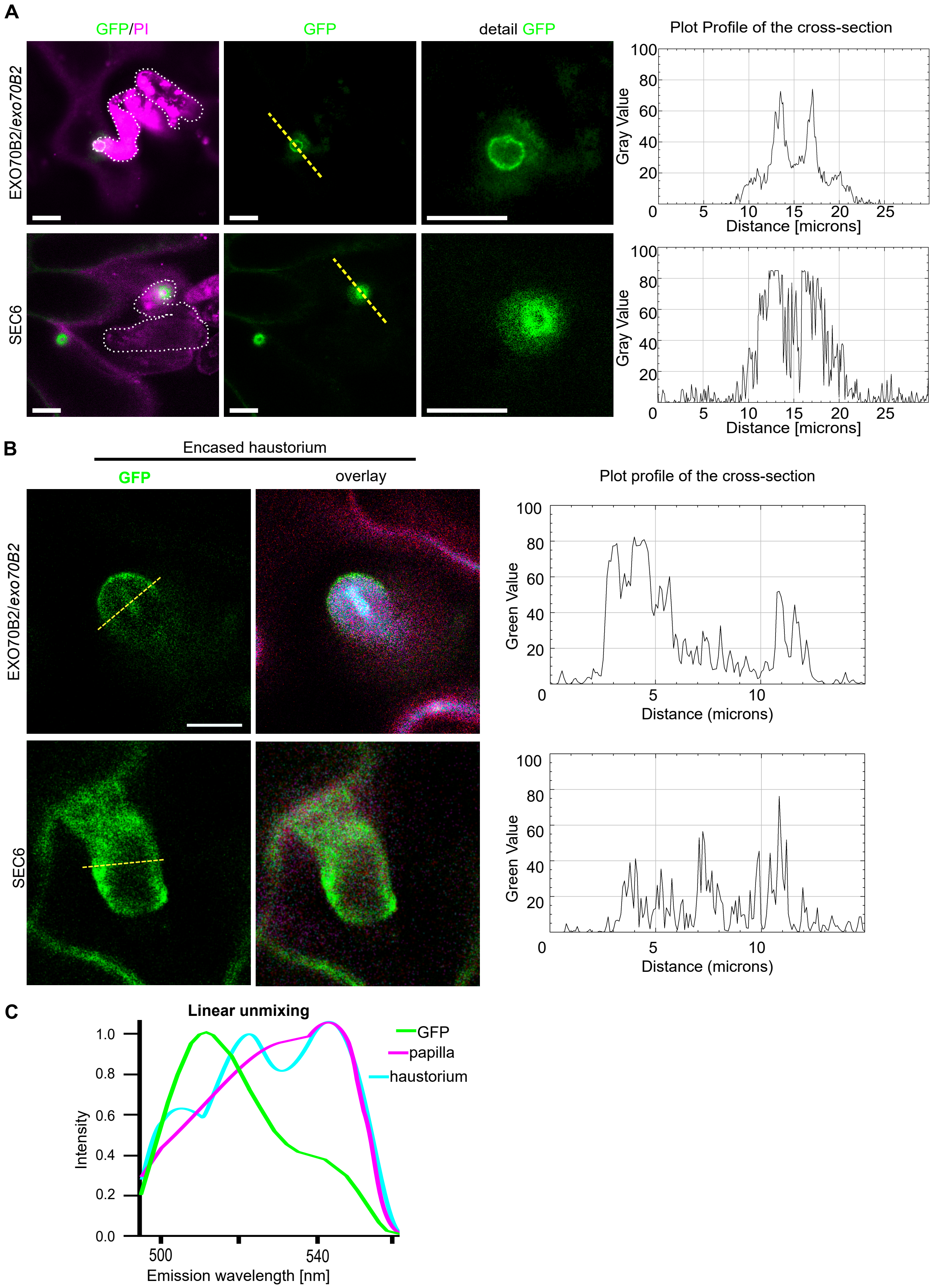


**Figure S3. Linear unmixing analysis of GFP-tagged exocyst subunits in papillae and haustoria.** (A) Focal accumulation of complemented EXO70B2-GFP and SEC6-GFP lines at Bg contact sites with Arabidopsis leaves. The fungus was stained with propidium iodide PI (2%) as shown in the magenta channel. All images display single plane sections obtained by confocal scanning microscopy with the same intensity of signal adjusted in FIJI. The GFP channel represents the signal sorted from GFP fluorophore specific emission spectra using the lambda scan mode of the microscope λ (505-511). The white dotted line highlights fungal structures. The yellow dashed line emphasizes the sections through papillae for signal profiling, as shown in the graph on the right. Scale bars represent 10 µm. (B) The accumulation of the GFP signal of complemented EXO70B2-GFP and SEC6-GFP after linear unmixing processing and the overlay shows the merged channels of pure GFP, non-specific signal of haustoria and papillae in the green channel. (C) Graphical representation of the pure GFP, unspecific papilla and haustorium spectrum used for linear unmixing analysis. The lambda scan was performed to visualize GFP alone with a 488 nm argon laser with a wavelength range from 489-600 nm for emission detection. The free-GFP plants were used for GFP spectrum characterisation. The non-transformed plants were used as the control for fungal structure spectrum characterisation.

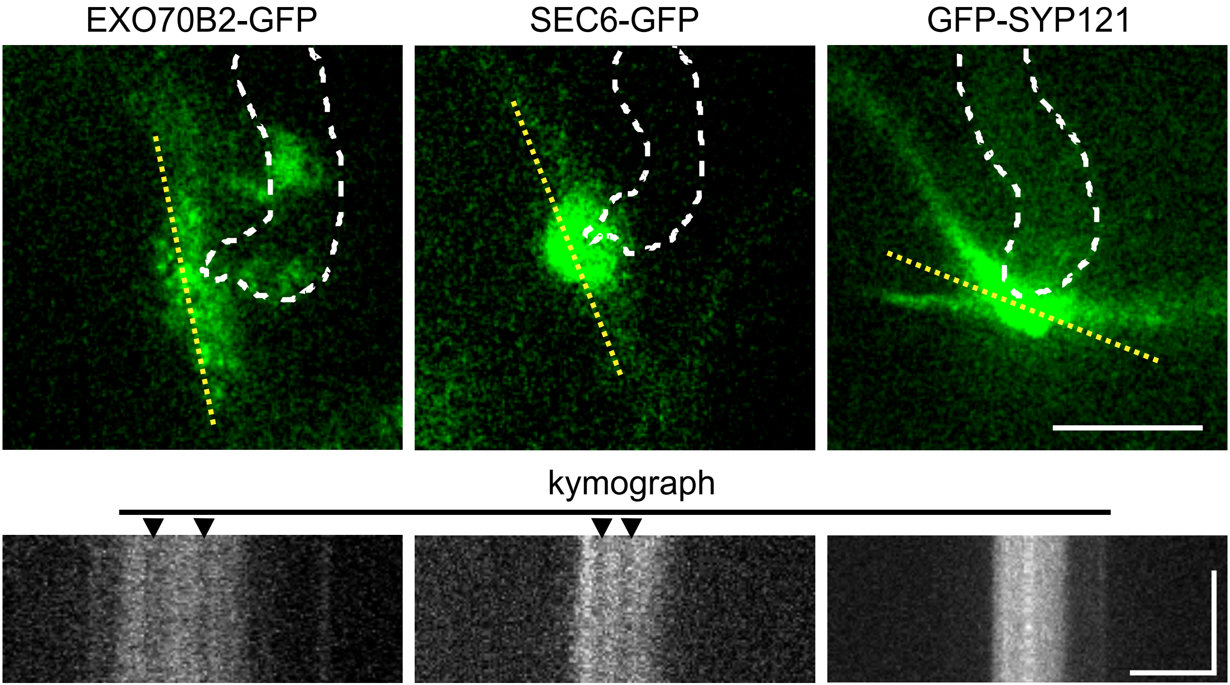


**Figure S4. Exocyst dynamics in papillary membrane domains.** The dynamics of GFP-tagged EXO70B2, SEC6 exocyst subunits and SYP121 SNRE in a papillary cross-section after Bg inoculation. The images on the left show the start section of the time series used for the kymographs (below); the scale bar represents 10 µm. The yellow dashed line defines the X-axis of the kymographs. All proteins were expressed under their natural promoters. For each kymograph, the vertical scale bar represents 30 s, and the horizontal scale bar is 3 µm. The white dashed line outlines the fungal structures. The black arrowheads point to gaps in the GFP signal in the papilla body. All images represent a single optical section obtained by confocal spinning disc microscopy.

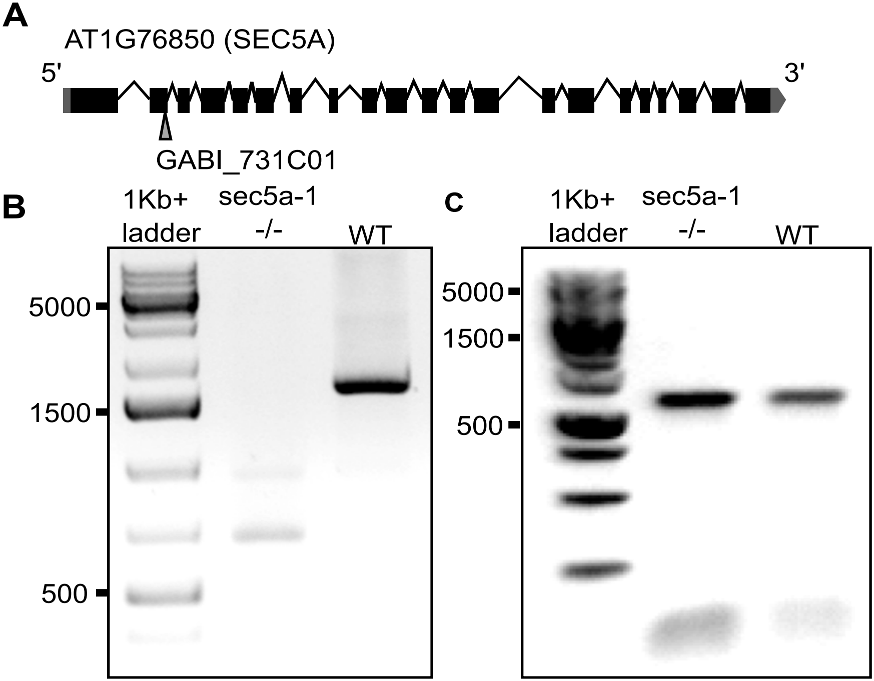


**Figure S5. Verification of the KO *sec5a-1* mutant.** (A) Graphical illustration representing the genomic sequence of the *Arabidopsis* SEC5A gene (B) The 1800-bp-long segment of SEC5A was amplified from the cDNA of 14-days-old *sec5-1* mutant and WT seedlings. (C) From the same cDNA samples, the 600-bp-long fragment of the control gene EXO70B2 was amplified as the cDNA quality control. The primers used for PCR were designed to cover the part with the insertion and to distinguish SEC5A from SEC5B (Supplementary Table 1).

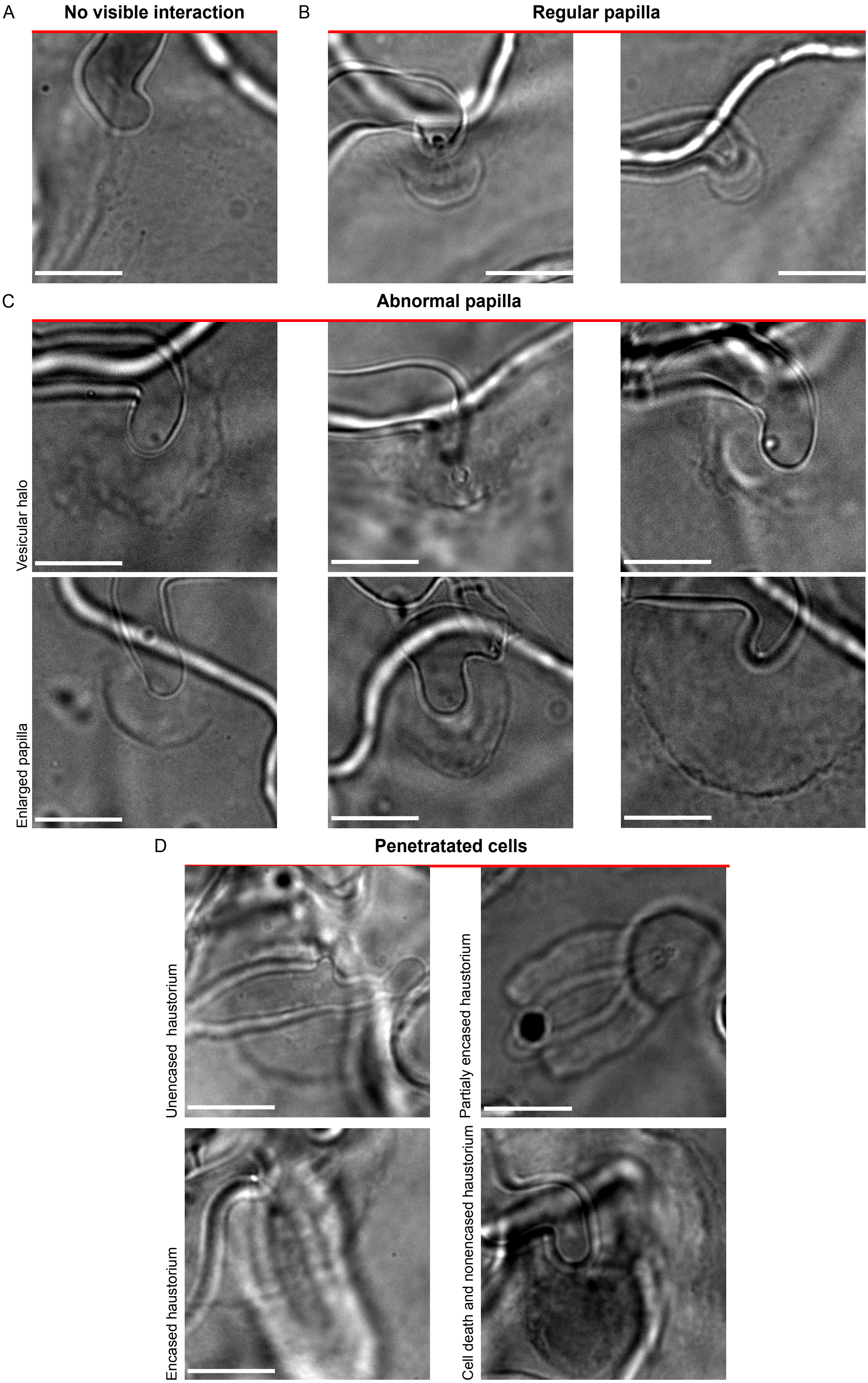


**Figure S6. Detailed visualisation of *Bg/*Arabidopsis interactions using trypan blue staining.** The images show structures spotted in a leaf of the exocyst mutants at 48 hpi with Bg. Leaves were stained with trypan blue, decolourized with chloral hydrate solution overnight and observed. All images represent a snapshot using the BW camera Axiom Widefield Zeiss Microscope with 100x objective. Observed interactions are categorized as follows: (A) no visible interaction, (B) regular papilla, (C) abnormal papilla, and (D) penetrated cells (with visible haustoria). Scale bars represent 10 μm.

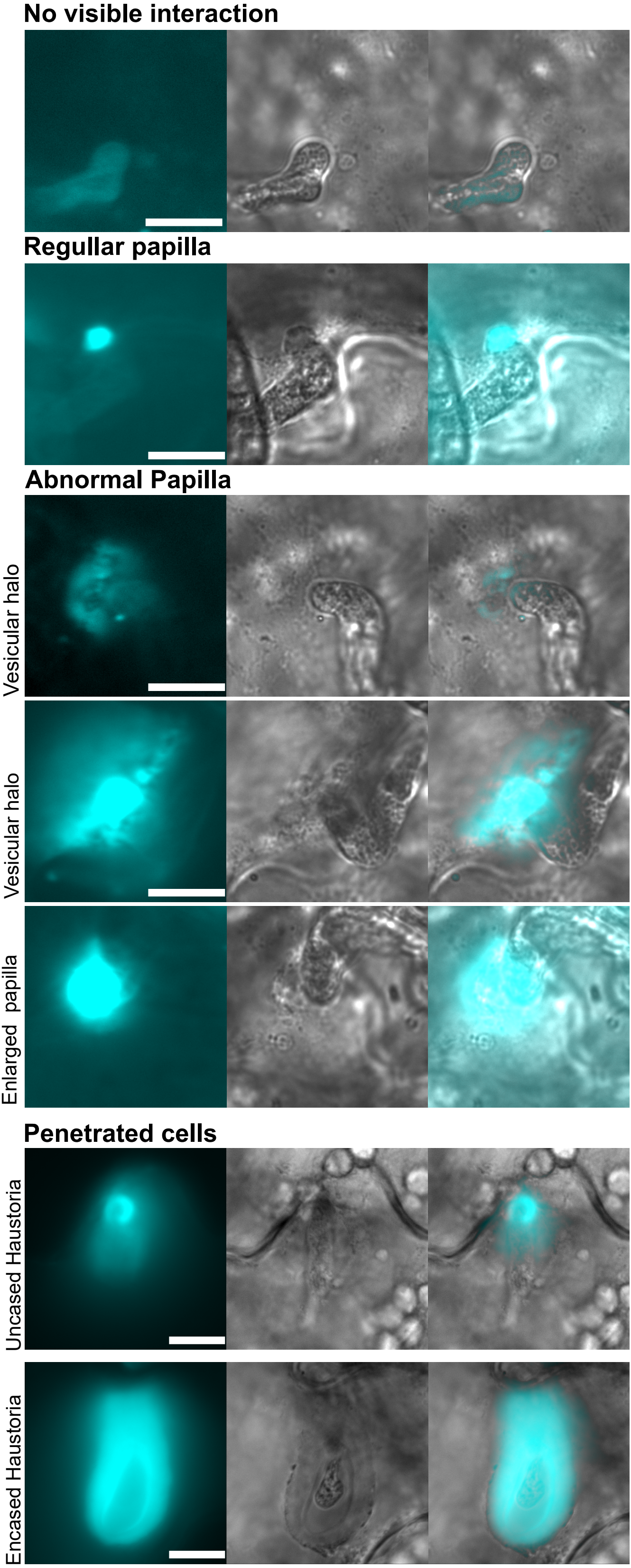


**Figure S7. Detailed visualisation of Bg/Arabidopsis interactions using aniline blue staining.** Four-weeks-old plants were treated with Bg spores for 24 h and then stained with aniline blue. Images in each panel from the left: callose aniline blue positive signal after UV excitation, brightfield and merged channel. All pictures represent a snapshot using the black/white camera Axiom Widefield Zeiss Microscope with a 100x objective. Scale bars represent 10 μm.

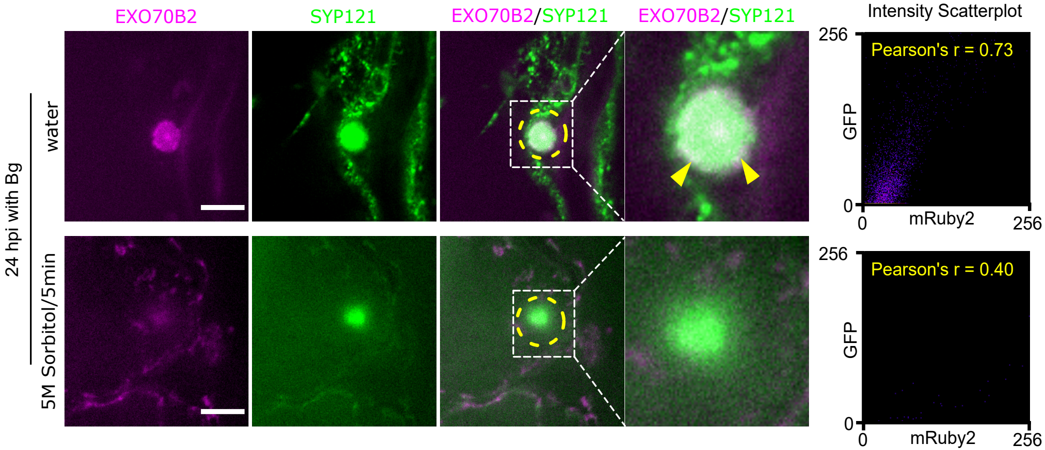


**Figure S8. EXO70B2 does not follow SYP121 into apoplast in defensive papilla.** Co-localisation of mRuby2-EXO70B2 and GFP-SYP121 in the membrane domain of a defensive papilla. The transgenic plants with the mRuby2-EXO70B2 and GFP-SYP121 were observed at 24 hpi with *Bg* in a normal water or hyperosmotic condition; images represent one of 15 analysed papillae with similar localisation. Pearson's correlation graph of mRuby2-EXO70B2 (Ch1) and GFP-SYP121 (Ch2) for non-plasmolysed and plasmolysed cells is shown on the right. The scale bar represents 5µm. The yellow dashed line marks the cross-section used for analysis. All images represent single plane sections.

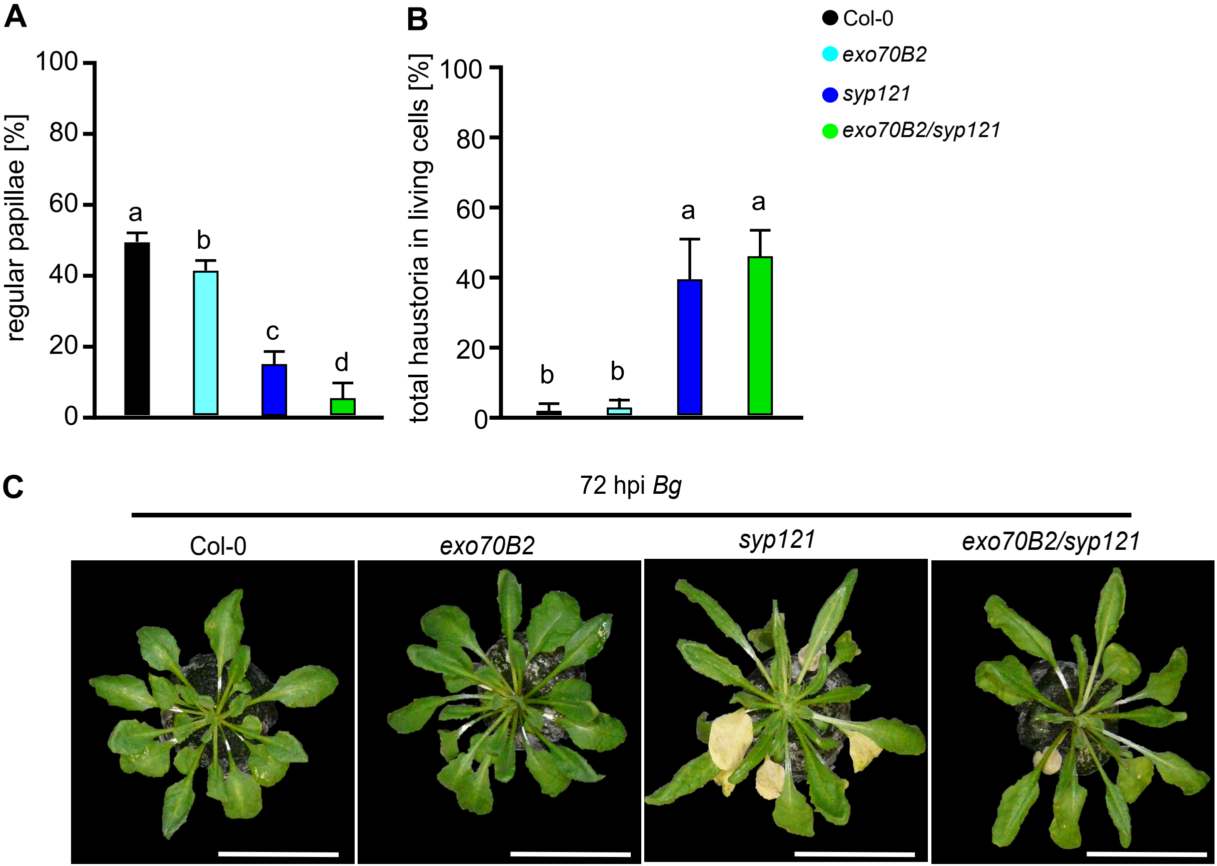


**Figure S9. Analysis of *exo70B2/syp121* double mutant reaction to Bg infection.** (A) Percentages of regular papillae from Bg spores per leaf in Col-0 and mutant lines *exo70B2, syp121, exo70B2/syp121,* 48 hpi. (B) Percentages of developed haustoria of Bg in epidermal leaf cells without visible cell death analysed for the same set of lines. n = 5. Small letters indicate significant differences analysed by ANOVA post-hoc Tukey-Kramer (HSD), Z score $p<0.01$. Error bars represent standard deviations. (C) Appearence of 4-weeks old Col-0, *exo70B2, syp121* and *exo70B2/syp121* plants 72 hpi with Bg. Scale bars represent 5 cm.

| Primers used for cloning | | | | | |
|---|---|---|---|---|---|
| **Name** | **Sequence** | **Gene of interest** | **Vector** | **Destination vector** | **Source fo used vectors** |
| **B2 prom EcoRI for** | AAGAATTCGAGCTCCGACGGACGAG | AtEXO70B2 | pENTR3C (Gateway) | pGWB4 | Nakagawa et al., 2007 |
| **B2 nostopXhol rev** | AAACTCGAGAAACTTGAGCTTTCCTTGAAC | | | | |
| **B2prom** | GGGGACAACTTTGTATAGAAAAGTTGCTGCCATTGGTATTGGTGG | AtEXO70B2 | pDONR P4-P1R (Gateway) | pB7m34GW | Karimi et al. 2005 |
| **B2prom rev** | GGGGACTGCTTTTTTGTACAAACTTGCATGATTGGATGGGAATTAATAATGT | | | | |
| **EXO70B2 CDS for** | GGGGACAAGTTTGTACAAAAAAGCAGGCTTAATGGCTGAAGCCGG | AtEXO70B2 | pDONR P2R-P3 (Gateway) | pB7m34GW | Karimi et al. 2005 |
| **EXO70B2 CDS rev** | GGGGACCACTTTGTACAAGAAAGCTGGGTAACTTGAGCTTTCCTTGAACA | | | | |
| **B1 prom for** | ATAGAAAAGTTGAATGCGGTAGAAGAGAGGATA | AtEXO70B1 | pDONR P4-P1R (Gateway) | pB7m34GW | Karimi et al. 2005 |
| **B1 prom rev** | TTGTACAAACTTGAGATTGAAACAGATGTGGAACC | | | | |
| **B1 CDS for** | CTTGTACAAAGTGGCTATGGCGGAGAATGGT | AtEXO70B1 | pDONR P2R-P3 (Gateway) | pB7m34GW | Karimi et al. 2005 |
| **B1 CDS rev** | GTATAATAAAGTTGTCATTTTCTTCCCGTGGTA | | | | |
| **B2 mRuby2 F** | AGTGAATTCTTCTCTGTTTATCCTCTCTATGC | AtEXO70B2 | TaqRFP-AS-N (Evrogen) | pBGWT | Karimi et al., 2002 modified by Sabol et al. 2017 |
| **B2 mRuby N rev** | TAACTCGAGCAACTTGAGCTTTCCTTGA | | | | |
| **EcoRVSec5a rev** | CTATAGTCTTCGTCTGGGTCGGG | AtSEC5a | pENTR3A (Gateway) | pUBvector | Grefen |
| **KpnISec5a for** | GGTACCgATGTCGAGCGATAGCAATG | | | | |
| **SYPdeltaC1Eco** | AAAGAATTCATGAACGATTTGTTTTC | AtSYP121 | pGADT7 (Clontech) | pGADT7 | Takara EU bio / http://www.clontech.com/US/Products/Protein_Interactions_and_Profiling/Yeast_Two-Hybrid/Vectors# |
| **SYPdeltaC1Sal** | AGTCGACTCATTTTCGCGTGTTCT | | | | |
| **GFP_For** | ATGGTGAGCAAGGGCG | eGFP | pDONR 221 (Gateway) | pB7m34GW | Karimi et al. 2005 |
| **GFP_Rev** | CTTGTAGTTGCCGTCGTCC | | | | |
| **mRUBY2 For** | ACCGGTAATGGTGTCTAAGGGCGAAGAG | mRUBY2 | TagRFP-AS-N (Evrogen) | pBGWT | Karimi et al., 2002 modified by Sabol et al. 2017 |
| **mRuby2 Rev** | TTTGCGGCCGCTTACTTGTACAGCTCGTCCATCC | | | | |

**Primers used for qPCR and semiqPCR**

| | |
|---|---|
| **qUBQ10 F** | GGCCTTGTATAATCCCTGATGAATAAG |
| **qUBQ10 R** | AAAGAGATAACAGGAACGGAAACATAGT |
| **qSEC8F** | GGGAATGGCGCCTTTCATCTCTGG |
| **qSEC8R** | GCTGCCATGGCCTGTTCCACTGC |
| **qEXO70B2F** | GAAGCACGCAGCGAAACTGAGGC |
| **qEXO70B2R** | GCACCTTACACACCTCATCGAACTGTG |
| **sqSEC5F** | TACAATCAGTGGAATTCCCCAGC |
| **sqSEC5R** | GTTGACTCTAATGGGGCTGA |
| **sqSEC15bF** | GCAATCGTCGAAAGGACGGC |
| **sqSEC15bR** | AGCCATACCTGCGTAGAC |
| **qACT7F** | GCCGATGGTGAGGATATTCAGC |

| | |
|---|---|
| **sqACT7R** | CAAACTCACCACCACGAACCAG |
| **sqEXO70BF** | GGCGGTGGGATTCACCCG |
| **sqEXO70B2R** | ACGCCTCCCATTAATCTCCG |
| | **Primers used for genotyping** |
| **SEC5aF** | CCTGTGGCTGCGGCTG |
| **SEC5aR** | CCTCCTCGAGTACACGCTTG |
| **GABI_08474** | ATAATAACGCTGCGGACATCTACATTTT |
| **SYP121CapsF** | CAACGAAACACTCTCTTCATGTCACGC |
| **SYP121CapsR** | CATCAATTTCTTCCTGAGAC |
| **digestion with MLU-1** | **100bp shift on gel** |
| **EXO70B2F** | CGTGATCCGTCTTTGTGTTTC |
| **EXO70B2R** | ACGCCTCCCATTAATCTCCG |
| **LB3** | CATCTGAATTTCATAACCAATCTCG |
| **EXO70B1F** | TTCGTTTATGGAGGTTTGTCG |
| **EXO70B1R** | TGGTCATTTAGCAGGTGGTTC |
| **GABI_08474** | ATAATAACGCTGCGGACATCTACATTTT |
| **SEC8R (Cole et al. 2005)** | CCTGCTTCTCCTTTATGATTTCACC |
| **SEC8F (Cole et al. 2005)** | CACGTAGGGAGGAGGGAATGG |
| **LBB sec8-4** | ATTTTGCCGATTTCGGAAC |
| **SEC15bF** | TTCACCAATAGCCAACTGACC |
| **SEC15bR** | ACTAAGGACATTTATACCTACCAACTG |
| **LBB** | ATTTTGCCGATTTCGGAAC |
| **EXO70A1F** | CTAGACGTTTGCAGCATCCTAT |
| **EXO70A1R** | ATATGTGTAATGCATTGGAGAAGC |
| **LBB** | ATTTTGCCGATTTCGGAAC |

List of proteins identified by immunoprecipitation of GFP tagged EXO70B2 after incubation with a pathogen.

| Accession | Protein | Gene name | Locus | Peptide counts |
|---|---|---|---|---|
| F4JZY1 | COP1-interactive protein 1 | CIP1 | At5g41790 | 9 |
| Q96318 | 12S seed storage protein | CRC | At4g28520 | 5 |
| P15456 | 12S seed storage protein | CRB | At1g03880 | 2 |
| Q39085 | Delta(24)-sterol reductase | DIM | At3g19820 | 5 |
| O80448 | Pyridoxal 5'-phosphate synthase subunit PDX1.1 | PDX11 | At2g38230 | 3 |
| Q9LMJ4 | Exocyst subunit EXO70B2 | EXO70B2 | At1g07000 | 4 |
| Q9C865 | SH3 domain-containing protein 1 | SH3P1 | At1g31440 | 2 |
| O64644 | Histone deacetylase complex subunit SAP18 | | At2g45640 | 2 |
| F4J8V9 | Actin 2 | ACT2 | At3g18780 | 2 |
| Q6NQ63 | protein belonging to family of DUF 3133 | AT1G01440 | At1g01440 | 2 |
| Q9LW76 | Ras-related protein RABG3c | RABG3C | At3g16100 | 2 |
| F4J8J7 | Uncharacterized protein | At3g11930 | At3g11930 | 1 |
| O04331 | Prohibitin-3 | PHB3 | At5g40770 | 1 |
| P39207 | Nucleoside diphosphate kinase 1 | NDK1 | At4g09320 | 1 |
| O80845 | Peroxisomal membrane protein 11D | PEX11D | At2g45740 | 1 |
| Q42290 | Probable mitochondrial-processing peptidase subunit beta, mitochondr | At3g02090 | At3g02090 | 1 |
| Q9S757 | Bifunctional L-3-cyanoalanine synthase/cysteine synthase C1, mitocho | CYSC1 | At3g61440 | 1 |
| F4JVC0 | Glycine-rich RNA-binding protein 8 | CCR1 | AT4G39260 | 1 |
| F4JJE5 | Putative proteasome subunit alpha type-4-B | PAC2 | At4g15165 | 1 |
| Q96262 | Plasma membrane-associated cation-binding protein 1 | PCAP1 | At4g20260 | 1 |
| F4KGV2 | 14-3-3-like protein GF14 lambda | GRF6 | At5g10450 | 1 |
| Q43729 | Peroxidase 57 | PER57 | At5g17820 | 1 |
| B3H4S6 | Dicarboxylate transporter 1 | DiT1 | At5g12860 | 1 |
| Q8LDS7 | B-cell receptor-associated 31-like protein | At1g48440 | At1g48440 | 1 |
| Q67YV9 | vesicle-associated membrane protein | VAMP721 | At1g04740 | 1 |
